# Computational identification of natural senotherapeutic compounds that mimic dasatinib based on gene expression data

**DOI:** 10.1101/2022.05.26.492763

**Authors:** Franziska Meiners, Riccardo Secci, Salem Sueto, Georg Fuellen, Israel Barrantes

**Affiliations:** Institute for Biostatistics and Informatics in Medicine and Aging Research, Rostock University Medical Center, Germany

**Keywords:** Drug Repurpositioning, Dasatinib, Piperlongumine, Senolytics, Gene Expression, Transcriptomics

## Abstract

The highest risk factor for chronic diseases is chronological age, and age-related chronic diseases account for the majority of deaths worldwide. Targeting senescent cells that accumulate in disease-related tissues presents a strategy to reduce disease burden and to increase healthspan.

Our goal was the computational identification of senotherapeutic repurposing candidates that potentially eliminate senescent cells, based on their similarity in gene expression effects to dasatinib, a tyrosine-kinase inhibitor that induces apoptosis in certain senescent cell types, and that is frequently used as a senolytic together with quercetin.

The natural senolytic piperlongumine (a compound found in *long pepper*), and the natural senomorphics parthenolide, phloretin and curcumin (found in various edible plants) were identified as potential substitutes of dasatinib. The gene expression changes underlying the repositioning highlight apoptosis-related genes and pathways. The four compounds, and in particular the top-runner piperlongumine, may be combined with quercetin to obtain natural formulas emulating the dasatinib + quercetin (D+Q) formula that is frequently used in clinical trials targeting senescent cells.

## 1. Introduction

Cellular senescence was investigated by Hayflick and Moorhead as early as 1961. They found that cultured human fibroblasts could only undergo a certain number of replications until a state of replicative arrest was entered, now known as replicative senescence (Campisi, 2000; Fyhrquist et al., 2013; Hayflick & Moorhead, 1961). Non-replicative cellular senescence can also be triggered by various factors including DNA damage, sustained inflammation, radiation, UVB light, DNA damaging chemotherapeutics, oncogene-activation, or PTEN tumor suppressor loss (Childs et al., 2014). Senescent cells feature cell cycle arrest, but they do not undergo apoptosis and instead remain metabolically active, usually displaying the so called senescence-associated secretory phenotype (SASP), the secretion of a diverse, often deleterious collection of pro-inflammatory cytokines, chemokines and proteases, leading to inflammation and tissue damage. Senescent cells accumulate with increasing age and contribute substantially to age-associated diseases. In senescent cells, signaling pathways are activated that sustain their resistance to apoptosis and, in contrast to proliferating cells, senescent cells are believed to need these pathways, termed senescent cell anti-apoptotic pathways, in order to stay alive (Y. Zhu et al., 2015).

As a consequence, a lot of effort has been invested into finding drugs (termed senolytics) that kill senescent cells, e.g., by inhibiting anti-apoptotic pathways. By disabling these pro-survival pathways, they enable the selective elimination of senescent cells via the induction of apoptosis (Y. Zhu et al., 2017). Target genes of senolytics include BCL2-family proteins such as BCL2L1, the kinases PIK3CA and AKT, the transcription factor TP53, cyclin-dependent kinase inhibitor 1A (CDKN1A, also known as p21), the chaperone protein HSP90, and plasminogen-activated inhibitor-2 (SERPINB2) (Zhu, et al., 2015; Zhu et al., 2020). Unlike senolytics, senomorphics are drugs that can suppress SASP factors or hinder stressed cells from becoming senescent, e.g. by activation of the NRF2 or FOXO pathways, by decreasing/inhibiting NF-κB or mTOR activity, by inhibition of IκB kinase (IKK), by scavenging free radicals, or by inhibiting the JAK-pathway (Martel et al., 2020; Niedernhofer & Robbins, 2018; Romashkan et al., 2021). We use the term “senotherapeutic” to cover senolytics and senomorphics.

Two known senolytics are dasatinib and quercetin, often studied in combination. Dasatinib is a drug developed for the treatment of leukemia, and it exerts its antitumoral activity by dual inhibition of SCR/ABL1 kinases, and by inhibiting the BCL-ABL1 fusion protein that causes chronic myeloid leukemia (Braun et al., 2020; Hochhaus & Kantarjian, 2013). Quercetin is a polyphenol (flavonol) known as a potent natural antioxidant, found in many fruit and vegetables (Bravo, 1998; Drewnowski & Gomez-Carneros, 2000). The joint senolytic activity of the two compounds was discovered through a hypothesis-driven approach (Kirkland & Tchkonia, 2020). Dasatinib can act as a senolytic through ephrin-dependent receptor ligands, partly by inhibition of SRC-kinase (Zhu et al., 2015; Kirkland and Tchkonia, 2020), and quercetin can act as senolytic partly by inhibition of the BCL2-family protein BCL2L1 and HIF1A, and PIK3CA (Zhu et al., 2015; Kirkland and Tchkonia, 2020).

Senolytics are cell-type specific (see Table 2); dasatinib and quercetin both target human preadipocytes and human umbilical vein endothelial cells (HUVECs) but with different effectivities, i.e. quercetin is more effective in killing HUVECs than preadipocytes, and dasatinib kills senescent human preadipocytes more effectively than HUVECs (Zhu et al., 2015). Combining dasatinib and quercetin (D+Q) successfully reduced viability in both cell types (Zhu et al., 2015). Further, D+Q reduced abundance of senescent primary mouse embryonic fibroblasts and senescent bone marrow derived mesenchymal stem cells (Zhu et al., 2015), and induced apoptosis in senescent (fibrotic) alveolar epithelial type II cells, as was shown in an ex vivo model of lung fibrosis (Lehmann et al., 2017). In murine models, D+Q prevented uterine age-related dysfunction and fibrosis, reduced intestinal senescence and inflammation and modulated the gut microbiome in aged mice, reduced senescent cell load in the context of age-related hepatic steatosis, and protected retinal ganglion cell loss by early removal of senescent cells (Cavalcante et al., 2020; Ogrodnik et al., 2017; Rocha et al., 2020; Saccon et al., 2021). Long-term treatment by D+Q reduced the number of senescent cells and ameliorated age-dependent intervertebral disk-degeneration in mice, along with downregulation of circulating proinflammatory factors and an increase in physical strength (Novais et al., 2021). In humans, D+Q showed improved disease-related outcomes e.g. leading to reduced adipose tissue senescent cell burden in individuals with diabetic kidney disease and it improved physical strength and function in patients with idiopathic pulmonary fibrosis (Hickson et al., 2019; Justice et al., 2019). However, as with other antitumor drugs, adverse events of dasatinib are frequent, such as respiratory events, skin irritation, myelosuppression, fluid retention events or diarrhea (Justice et al., 2019). Hence, finding non-toxic analogs of dasatinib, especially from natural sources, possibly for combination with quercetin, would be of high value.

In this regard, traditional drug discovery is a time-consuming, costly and labor-intense process with high failure rates (Everett, 2015). Computational methods to identify existing drugs for a new purpose (known as drug repositioning, or repurposing) offers an alternative to de novo drug discovery, as it imposes fewer risks, resources and economic effort (Jarada et al., 2020; Lima et al., 2019; Xue et al., 2018). A common repositioning approach is built on the hypothesis that if two drugs induce similar gene expression profiles and thus may be assumed to have similar modes of action, both could be considered to treat the same condition (Jarada et al., 2020). Transcriptomic gene expression profiles capture some of the dynamics of the cellular response to a drug intervention and measure the transcriptional activity of hundreds or thousands of genes simultaneously, and therefore help understanding how genes act under the same or similar circumstances (Jarada et al., 2020). Two key resources for drug repositioning are the Connectivity Map and the Library of Integrated Network-Based Cellular Signatures (LINCS) projects. The Connectivity Map is a database where genes, drugs and diseases are connected by common gene expression signatures (Subramanian et al., 2017). LINCS, in turn, is a program funded by the National Institutes of Health to generate an extensive reference database of cell-based perturbation-response signatures (Koleti et al., 2018). LINCS is an expanded version of the Connectivity Map and comprises over a million gene expression profiles of chemically perturbed human cell lines that can be used to discover mechanisms of action of small molecules, based on a compacted representation of the transcriptome (Duan et al., 2016; Subramanian et al., 2017).

Because the dasatinib and quercetin (D+Q) combination has been studied extensively in regard to cellular senescence and senolysis, and gene expression data of dasatinib-interventions are available online, the gene expression-based approach of repositioning was used to find candidate compounds that may replace dasatinib, to be used together with quercetin as a senolytic combination. Therefore, here we aimed to identify compounds that show similar senolytic activity as dasatinib through computational drug repositioning, focussing on compounds found in dietary sources that could act as safe substitutes of dasatinib. More specifically, we aimed to (i) find studies about dasatinib that include publicly available gene expression data; (ii) identify differentially expressed genes (DEGs) associated to senescence and aging in these dasatinib intervention studies; (iii) search for dasatinib analogs, especially natural compounds, based on the DEGs related to dasatinib, employing the LINCS data and (iv) use the gene expression data underlying the repositioning to find hypotheses for potential senotherapeutic molecular mechanisms that dasatinib and its analogs may have in common. The molecular-mechanistic insights from (ii) and (iv) suggest that the gene expression profile of dasatinib that we used for the repositioning is strongly linked to cellular senescence and apoptosis, as are the gene expression changes underlying the repositioning in case of the analogs (specifically, in case of piperlongumine). Thus, our approach should give us maximum confidence in senotherapeutic effects, also *in vivo* in humans, by the analog itself or, at least, by the analog in combination with quercetin.

## 2. Results

We considered gene expression data from the Gene Expression Omnibus (GEO) (Clough & Barrett, 2016) describing (1) the long-term effects of dasatinib in the AML cell line Kasumi-1, (2) transcriptomic differences in dasatinib-sensitive and dasatinib-resistant prostatic cancer cell lines, (3) the effect of dasatinib-treatment on the breast cancer cell line MDA-MB-468 (see Table 1).

**Table 1:**
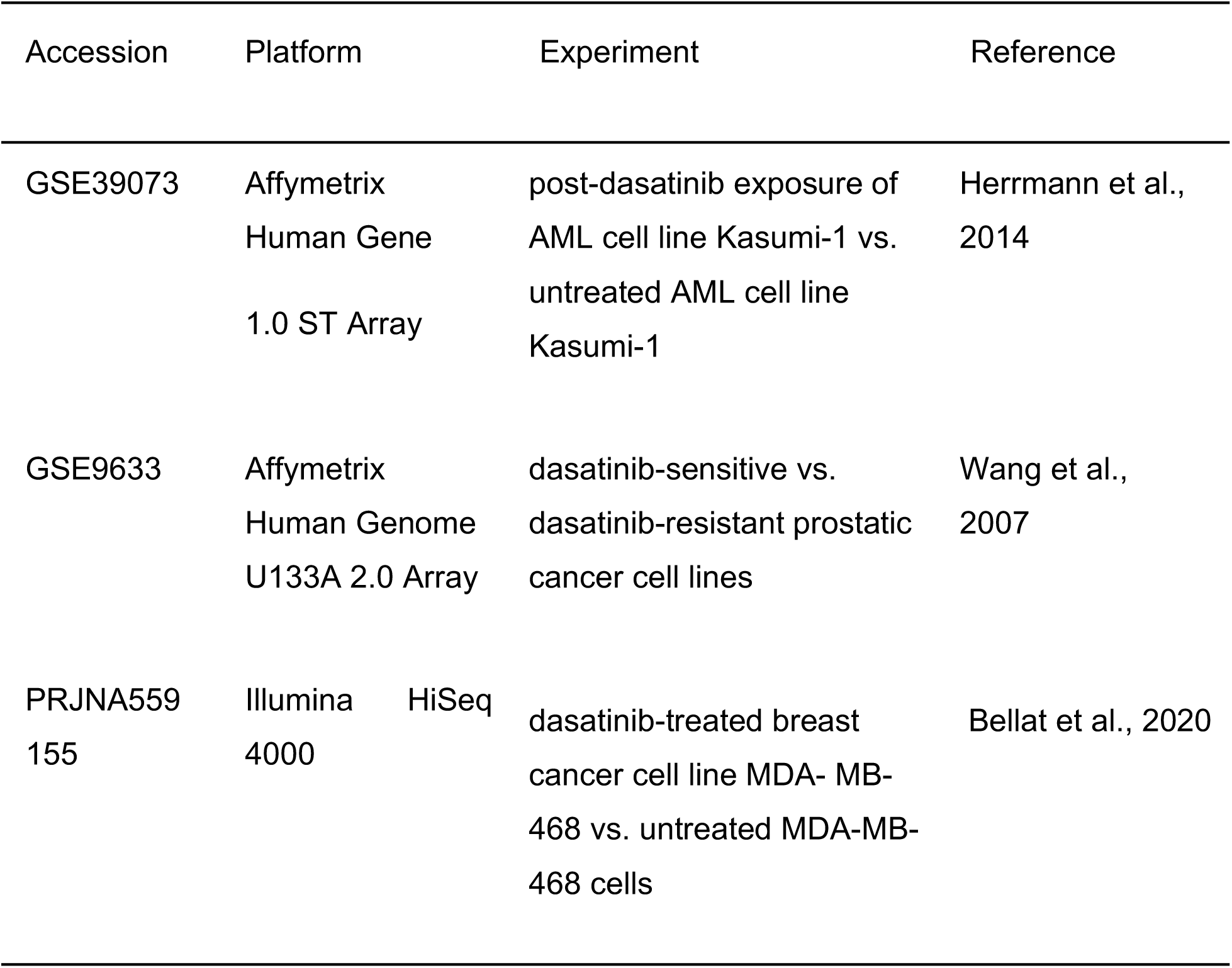
List of datasets used in this study. AML: Acute myeloid leukemia

**Table 2:** The known senolytic compounds Dasatinib (D), Quercetin (Q), the combination of both (D+Q), and Piperlongumine, and the targeted senescent cell types

| Compound | Targeted senescent cell types | Reference |
| --- | --- | --- |
| Piperlongumine | human WI fibroblasts | (Y. Wang et al., 2016) |
| Dasatinib (D) | human and mouse preadipocytes<br>HUVECs | (Y. Zhu et al., 2015b) |
| Quercetin (Q) | HUVECs<br>human and mouse preadipocytes | (Y. Zhu et al., 2015b) |
| D+Q | same cells as by D or Q<br>primary mouse embryonic fibroblasts<br>bone marrow-derived mesenchymal stem cell<br>alveolar epithelial type II cells<br>reduction in senescent cells in skin ulcer samples | (Gasek et al., 2021; Lehmann et al., 2017; H. Wang et al., 2020; Zhou et al., 2021; Y. Zhu, et al., 2015b) |

### 2.1 Genes associated with aging and cellular senescence, and with biological processes associated with apoptosis, in the treated Kasumi-1 (AML) cell line

GEO accession GSE39073 entailed microarray gene expression profiles from AML-derived Kasumi-1 cells upon treatment with dasatinib. The aim of these experiments was to study the effect of the longterm exposure to dasatinib in leukemic cells, which usually triggers drug resistance and thus is a major problem for the treatment of patients with AML (Herrmann et al. 2014). Here, the expression data from these experiments was re-analyzed to identify DEGs, which produced 190 up- and 192 downregulated genes between both conditions (Supplementary Table 1). From these DEGs, eight genes (KNYU, c-FOS, ITGB2, PRKCD, BCL2, MPO, APP, TIMP2) were annotated with the GO term aging and one gene, PRKCD, with the term cellular senescence (Supplementary Table 2). **PRKCD** (Protein kinase C) was upregulated with a log2 fold change (LFC) of 3.37 and it is a tumor suppressor protein and positive regulator of cell cycle progression; PRKCD may regulate apoptosis (see <u>NCBI Gene ID 5580</u>), and it plays a role in the regulation of senescence-induction in human diploid cells (Katakura et al., 2009). Also associated with cellular senescence, based on the literature, is the apoptosis regulator **BCL2**, which was upregulated with LFC=2.62. BCL2 is an integral mitochondrial membrane protein that blocks apoptosis of e.g. lymphocytes (<u>NCBI Gene ID 596</u>). It is a pro-survival protein and a target of senolytics inducing apoptosis, and may influence human lifespan (Ukraintseva et al., 2021; M. Zhu et al., 2020). Biological processes related to apoptosis were also enriched in the DEG list (adjusted p-value < 0.05) and included *cell death, programmed cell death, regulation of cell death, apoptotic process*, and *regulation of programmed cell death* (see Supplementary excel sheet “GO-gprofiler.5-23-22_AML”, for the enriched genes in the “intersections” column).

### 2.2 Genes associated with aging and cellular senescence, and with biological processes associated with apoptosis, in the dasatinib-sensitive prostatic cancer cell lines

GEO-Accession GSE9633 features base-line gene expression profiles of dasatinib-sensitive and dasatinib-resistant prostatic cancer cell lines. We identified 198 differentially expressed genes, with 138 upregulated and 51 downregulated genes between dasatinib-sensitive and dasatinib-resistant prostatic cancer cell lines. A large number of genes were annotated to the term *aging* (Supplementary Table 3), but some of these were also associated with cellular senescence in the literature, including **SERPINB5** among the upregulated genes, a tumor suppressor and senescence-associated marker (Bascones-Martínez et al., 2012; Sheng et al., 1996), the expression of which is linked to genotoxic and oxidative stress (Bianchi-Frias et al., 2010). **TGFBR2** was also upregulated in dasatinib-sensitive cell lines. This growth factor receptor may play a role in the interplay between cell survival and apoptosis in determining human lifespan (Ukraintseva et al., 2021) as it is involved in the phosphorylation of transcription factors associated with proliferation, cell cycle arrest, immunosuppression and tumorigenesis (<u>NCBI Gene ID 7048</u>). Another upregulated gene is **CDKN2A** (p16) that encodes a well-established marker of cellular senescence (Bernard et al., 2020). Accumulation of p16-positive cells (suggested to be senescent) during adulthood negatively influences lifespan and promotes age-dependent changes and diseases in various organs and tissues (D. J. Baker et al., 2016). Although the GO term “cellular senescence” was not enriched, the enrichment analysis showed that the cellular senescence pathway (KEGG accession ko04218) was enriched, and genes associated with this pathway were all upregulated, including some of the ones mentioned above (TGFBR2, TGFB2, HLA-A, CDKN2A, ZFP36L1, HLA-E, HLA-G, GADD45A, FOXO1, RRAS and GADD45B). Enriched biological processes also included processes associated with apoptosis (see Supplementary excel sheet “GO-gprofiler.5-23-22_PC-cancer” for enriched genes found in the “intersections” column), including *apoptotic process and positive regulation of apoptosis, programmed cell death* and *positive regulation of programmed cell death* (adjusted p-value < 0.05).

### 2.3 Genes associated with aging and cellular senescence, and with biological processes associated with apoptosis, in the MDA-MB-468 breast cancer cell lines

The gene expression dataset with the accession PRJNA559155 includes expression profiles of the dasatinib-treated breast cancer cell line MDA-MB-468. Differential expression analysis resulted in 189 upregulated and 80 downregulated genes between dasatinib-treated- and control MDA-MB-468 cells. Among the differentially expressed genes, some of the genes were annotated with the term *aging* (see Supplementary Table 4). Among these genes, the **CCL11** gene was most significantly downregulated with an LFC of -9.78 in the dasatinib-treated cell lines. CCL11 (also known as eotaxin-1) is considered to be an aging- and inflammation-associated plasma chemokine and a SASP-factor (Camell et al., 2021; Cameron et al., 2016). It acts as an eosinophil chemoattractant, is associated with allergic responses and Th2 inflammatory disease, colon tumorigenesis (Polosukhina et al., 2021), and with cell migration in rheumatoid arthritis (Wakabayashi et al., 2021). CCL11 is a putative biomarker for the prediction of severity and mortality of elderly patients with sepsis-induced myocardial injury (Li et al., 2020). The GO term “cellular senescence” was not enriched, and none of the GO biological processes were associated with apoptosis (see Supplementary excel sheet “GO-gprofiler_5-23-22_BC”).

### 2.4 Natural compound identification

Candidate compounds were identified using the L1000CDS^2^ webtool, using the output of 50 predictions (corresponding to LINCS perturbations) that mimic or reverse the input signature (the up- and downregulated genes identified from the differential expression analysis of dasatinib). *Reverse* matching was chosen for AML, and prostatic cancer (PC) datasets to reverse the disease-associated signature, so that input downregulated genes are intersected with input upregulated genes, and vice versa (Duan et al., 2016). *Mimic* was chosen for the breast cancer (BC) dataset to mimic the effect of dasatinib, intersecting downregulated genes with downregulated genes from the reference L1000 genes (and upregulated genes with upregulated genes). The selected compounds obtained from L1000CDS^2^ were labeled manually as *natural compounds* as appropriate (Table 3). In *reverse* mode, natural compounds from the AML-dataset were piperlongumine, parthenolide and curcumin on ranks 1, 20 and 40, respectively. Also in *reverse* mode, natural compounds from the PC-dataset were piperlongumine and parthenolide on ranks 27 and 39. In *mimic* mode, natural compounds from the BC-dataset were phloretin and parthenolide on ranks 7 and 32.

**Table 3:** Selected natural compounds mimicking the treatment with dasatinib. These compounds were identified using the L1000CDS^2^ tool. The rank is based on the overlap; the overlap is the score, based on the intersection length between the input DEGs and the signature DEGs divided by the effective input, i.e. the intersection-length between input genes and L1000 genes

| dataset GSE39073 – Kasumi-1 cells, *reverse* |  |  |  |  |
| --- | --- | --- | --- | --- |
| Rank | Score | Compound | Class of Compound | Source |
| 1 | 0.0634 | piperlongumine | amide alkaloid | *Piper longum* |
| 20 | 0.0387 | parthenolide | sesquiterpene lactone | *Tanacetum parthenium* |
| 40 | 0.0352 | curcumin | diarylheptanoid | *Curcuma longa* |
| dataset GSE9633 – prostatic cancer cell lines, *reverse* |  |  |  |  |
| Rank | Score | Compound | Class of Compound | Source |
| 27 | 0.071 | piperlongumine | amide alkaloid | *Piper longum* |
| 39 | 0.0656 | parthenolide | sesquiterpene lactone | *Tanacetum parthenium* |
| dataset PRJNA559155 – breast cancer cells, *mimic* |  |  |  |  |
| Rank | Score | Compound | Class of Compound | Source |
| 7 | 0.0435 | phloretin | dihydrochalcones | e.g. apples |
| 32 | 0.0348 | parthenolide | sesquiterpene lactone | *Tanacetum parthenium* |

Piperlongumine was the highest-ranking compound identified with the AML-dataset GSE39073. The highest overlap (in terms of overlapping genes) was seen with piperlongumine-treated NOMO1 cells (Duan et al., 2016); treatment dose was 10µm. NOMO1 is an AML cell line (Quentmeier et al., 2004) just like the Kasumi-1 cell line. Overlapping genes of the *input upregulated* and the piperlongumine-based *signature downregulated* genes were ACSL1, ATP8B4, CTSG, EIF1AY, FLT3, HCK, KDM5D, LYZ, PLAC8, PRKCD, PTPN6, RNASE2, RPS6KA1, TNFRSF10B and TNS3. Overlapping genes of the *input downregulated* and the *signature upregulated* genes were DDAH1, FBXO21, SLC38A1 and TSPAN13 (Supplementary Table 5). Enriched biological processes (adjusted p-value < 0.05) in the aggregate list of 19 genes include apoptosis-related processes (see Supplementary Table 6), e.g. *negative regulation of apoptotic process, intrinsic apoptotic signaling pathway, TRAIL-activated apoptotic signaling pathway, negative regulation of glial cell apoptotic process, negative regulation by symbiont of host apoptotic process* and *intrinsic apoptotic signaling pathway in response to oxidative stress*.

Piperlongumine was also found on rank 27 with the PC-dataset where PC3 cells were treated with 10µM piperlongumine. PC3 is a dasatinib-resistant prostate cancer cell line (Wang et al., 2007). *Input upregulated and signature downregulated* overlapping genes include AHNAK2, ALDH1A3, AREG, C3, CAPG, CST6, DDX60, FERMT1, ITGA3, KRT7, LAMA3, RAC2, RRAS, S100A2, TGFBR2 and ZBED2; one overlapping gene between *input downregulated and signature upregulated* genes was identified, which was LEF1 (see Supplementary Table 7). Here, the biological process *positive regulation of apoptotic cell clearance* was associated with the gene C3, and the gene RAC2 was associated with the process *engulfment of apoptotic cell clearance* (See Supplementary Table 8).

**Curcumin** was also identified from the AML dataset, on rank 40, where PL21 cells were treated with 48µM curcumin (Duan et al., 2016). PL21 is also an AML cell line (Kubonishi et al., 1984). Overlapping genes of the *input upregulated* and the curcumin *signature downregulated* genes were ADCY7, AHNAK, ATP8B4, BEX1, CXCR4, DDX3Y, EIF1AY, EPB41L3, KDM5D, PRKCD, RASSF2 and VIM. One gene overlapped with the *input downregulated* genes and the *signature upregulated* genes, which was MEST. Enriched apoptosis-associated biological processes (p-value < 0.05) included *regulation of glial cell apoptotic process* and *intrinsic apoptotic signaling pathway in response to oxidative stress*.

**Parthenolide** is a sesquiterpene lactone of the chemical class of terpenoids (Gali-Muhtasib et al., 2015) and was identified with L1000 using all three datasets, though at low ranks in all of these: on rank 20 with the AML-dataset, on rank 39 with the PC-dataset, and on rank 32 with the BC-dataset (Table 5). **Phloretin** is a dihydrochalcone flavonoid found in fruits such as apples, kumquat, pear, strawberry and in vegetables. Phloretin appeared on rank 7 from the expression signature of BC-dataset PRJNA559155.

## 3. Discussion

The process of aging involves most (if not all) aspects of life and in molecular terms, it thus involves a wide variety of signaling pathways at least to some degree. Aging is considered to be the causal process underlying age-associated disease and dysfunction (Fuellen et al., 2019). Accordingly, increasing chronological age is the foremost risk factor for the development of all kinds of chronic diseases such as type 2 diabetes, cardiovascular disease, osteoporosis, arthritis, Alzheimer’s disease and cancer, along with age-associated dysfunction such as frailty and sarcopenia. Shared molecular mechanisms behind these diseases and dysfunctions, and thus considered to be hallmarks of aging, include the accumulation of senescent cells, the buildup of macromolecular and genetic damage, metabolic dysfunction, loss of proteostasis and defective stem cell function (López-Otín et al., 2013).

Senescent cells contribute to aging and age-associated disease and dysfunction partly because of their high metabolic activity despite growth arrest, associated with the secretion of a complex, multi-component SASP which acts on the tissue microenvironment, usually in an unfavorable way (Wiley & Campisi, 2021). Resistance to apoptosis is a hallmark of senescent cells, primarily facilitated through upregulation of BCL2 family proteins; resistance to oxidative stress is another factor (Childs et al., 2014; X. Zhang et al., 2018). The SASP collection of proinflammatory cytokines, chemokines, bioactive lipids and damage-associated molecular patterns contribute to what is termed “inflammaging”, a chronic inflammation that is a common attribute in aged tissues that is – at least in part – due to the accumulation of senescent cells (Cevenini et al., 2013; Franceschi & Campisi, 2014; Wiley & Campisi, 2021). This accumulation leads to the abnormal activation of pathways such as NF-κB, that are needed to maintain many physiological functions, but when constitutively activated lead to accelerated aging (Amiri & Richmond, 2005; R. G. Baker et al., 2011; García-García et al., 2021; Salminen et al., 2008; L. Zhang et al., 2021). Still, the accumulation of senescent cells can be subject to “senotherapeutic” intervention: by direct killing (senolysis), by modification of the SASP (senomorphics) or simply by slowing down the process by which cells become senescent (gerostatics).

Using L1000CDS^2^ we obtained lists of compounds that have either similar or opposite gene expression profiles as compared to the input gene lists describing the action of dasatinib. Natural compounds were curated manually,, and we found four natural candidate-compounds as analogs of dasatinib, all of which are found in common foods and all of which have been under investigation already for their anti-inflammatory, anti-cancer or anti-aging effects: piperlongumine, phloretin, curcumin and parthenolide.

**Piperlongumine** is a known natural senolytic compound that was found based on the differentially expressed genes between dasatinib-treated Kasumi-1 (AML-dataset GSE39073) and untreated cells, which show an overlap with the L1000 dataset of the piperlongumine-treated AML cell line NOMO1. This overlap of DEGs featured an enrichment in genes and processes involved in apoptosis, including *positive regulation of apoptosis* and *programmed cell death*. Moreover, in the overlap with the PC-dataset, genes such as SERPINB5 and CDKN2A were differentially expressed, both encoding for senescence-associated markers. Piperlongumine is one of the few natural compounds shown to selectively kill senescent cells, that is, human WI-38 fibroblasts made senescent by ionizing radiation, replicative exhaustion or by expression of the oncogene Ras (Y. Wang et al., 2016; Y. Zhu et al., 2015) and therefore, it is a promising repurposing candidate in our context.

Piperlongumine is a natural compound with a strong safety record, and it has selective toxicity toward cancer cells and senescent cells, but does not induce significant toxicity in non-senescent, non-cancerous cells (Adams et al., 2012; Y. Wang et al., 2016), including peripheral blood T cells (PBTs) (Liang et al., 2020). 72h after incubating senescent WI human fibroblasts with piperlongumine leaves 30% of the senescent cells viable (Y. Wang et al., 2016), by the 10µM dose-regimen that was also used for the L1000 data. When combining piperlongumine with ABT-263 (navitoclax, a potent BCL2 inhibitor), a synergistic effect was observed, killing almost all senescent cells; the authors suggested that piperlongumine eradicated the subpopulation of senescent cells that was resistant to ABT-263 (Y. Wang et al., 2016). While BCL2 family proteins are thought to be primarily responsible for a senescent cells ability to resist apoptosis, and BCL2/BCL2L1/BCL2L2 inhibitors are effective senolytic drugs (e.g. ABT-263) (Chang et al., 2016), there is a concern that BCL2 inhibitors have on-target and off-target toxicities, such as thrombocytopenia and neutropenia (Rudin et al., 2012).

Data on piperlongumine’s mode of action in general, and specifically on how it induces apoptosis in cancer cells is available from a number of studies (e.g. Thongsom et al., 2017). Senescent cells and cancer cells share some pro-survival pathways and have in common e.g. active DNA damage responses (Ghosal & Chen, 2013), high metabolic activity including increased glycolysis (Dörr et al., 2013), and the reliance on dependence receptors to resist apoptosis (Goldschneider & Mehlen, 2010). Data from these studies, including the data we found from L1000 (especially the overlapping genes associated with apoptosis) based on repurposing dasatinib-associated expression data, thus suggest piperlongumine-induced apoptosis in senescent cells (Y. Wang et al., 2016; Y. Zhu et al., 2015a).

In more detail, piperlongumine has been shown to kill senescent fibroblasts without the induction of reactive oxygen species (Y. Wang et al., 2016), though it was later demonstrated that it inhibits the OXR1 (oxidation-resistance 1) protein that in turn leads to the expression of antioxidant genes. OXR1 is upregulated in senescent human WI38 fibroblasts and thus it is a proposed senolytic target (X. Zhang et al., 2018). When piperlongumine binds to OXR1 (see Supplementary Figure 1), the protein is degraded, leading to increased production of reactive oxygen species in senescent cells, mediated by low or zero levels of antioxidant genes such as heme oxygenase 1 (HMOX1), glutathione peroxidase 2 (GPX2) and catalase (CAT), presumably due to missing/reduced OXR1. Then, senescent cells are more susceptible to oxidative stress, leading to their apoptosis (X. Zhang et al., 2018; Bago et al., 2021). Of note, GPX2 (or glutathione) is the major hydrogen peroxide and organic hydroperoxide scavenger (also regulated by NRF2), induced by e.g. cigarette smoke (Singh et al., 2006).

Piperlongumine has also shown to interfere with T-cell differentiation and is considered to be a selective immunosuppressant (Liang et al., 2020), partly, again by a pro-oxidative action, here due to intracellular depletion of glutathione levels (Bago et al., 2021). This was linked to the inhibition of the transcription factors RORC (RORγt), HIF1A and STAT3, resulting in lowered production of IL22, IL17A, IL17F, and subsequent inhibition of Th17-differentiation, but not of regulatory Th1 and Th2 cells (Tregs), along with reduced expression of CD69 and CD35 expression markers (Bago et al., 2021; Liang et al., 2018, 2020). This is interesting and important, because the Th17/Treg ratio increases during aging, and increasing Th17/Treg imbalance possibly contributes to an altered pro-inflammatory/ anti-inflammatory immune response and thus indicates a higher risk to develop inflammatory diseases with increasing age (Schmitt et al., 2013).

In our analyses, we specifically focused on overlapping genes between the dasatinib-associated gene expression changes and piperlongumine-treated cells from the L1000 database, looking for apoptosis-related genes. One of the downregulated genes in the piperlongumine-based signature (Supplementary Table 5), PTPN6 (also known as SHP-1) is a tyrosine phosphatase that has been shown to interfere with cellular senescence via p16 signaling, and was proposed to regulate senescence in nasopharyngeal carcinoma (NPC) cells (Sun et al., 2015). Two downregulated overlapping genes were FLT3 and HCK, both enriched in the biological process *apoptotic process*, and additionally in the pathway *FLT3 signaling through SRC family kinases* (HAS-9706374). FLT3 and HCK are described as attractive targets for cancer therapy. Experimentally, FLT3 inhibition led to apoptosis in FLT3 positive AML cells (Lee et al., 2018), and dasatinib was shown to reverse induced resistance to FLT3-inhibition in the treatment of AML (Weisberg et al., 2012). HCK has an important role in the production of TNF and IL-6, enhances the secretion of growth factors, and targeting HCK has been proposed to alleviate excessive inflammation (Poh et al., 2015; Smolinska et al., 2011).

In the overlapping genes between the PC-data and the piperlongumine effects as known from L1000, the CS3 gene was positively associated with the regulation of apoptotic cell clearance (see supplementary excel sheet overlap_PL_PC3_PC-dataset), and it is downregulated in response to piperlongumine in PC cells.

Other identified compounds were phloretin, parthenolide and curcumin, which are described in more detail in the supplementary information.

To conclude, datasets corresponding to three different experiments studying the effects of dasatinib in gene expression were analyzed, from which we identified four natural compounds with potential senotherapeutic properties, all of which are readily available from dietary sources: the senolytic piperlongumine, and the senomorphics parthenolide, curcumin and phloretin.

The use of piperlongumine was described in cancer research, yet it would be interesting to investigate the systemic effect of piperlongumine (in terms of e.g. inflammatory markers in the blood) and its effect on overall health in humans. In particular, its combination with quercetin (“P+Q”) may be a natural-compound alternative to the combination of dasatinib and quercetin (“D+Q”) that was used by Hickson et al. (2018) and Justice et al. (2019) in the context of diabetic kidney disease and idiopathic pulmonary fibrosis; and is used in followup work, including a variety of senotherapy trials all over the world, see the clinicaltrials.org website.

## 4. Methods

See Figure 1 for a Methods overview.

**Figure 1.**
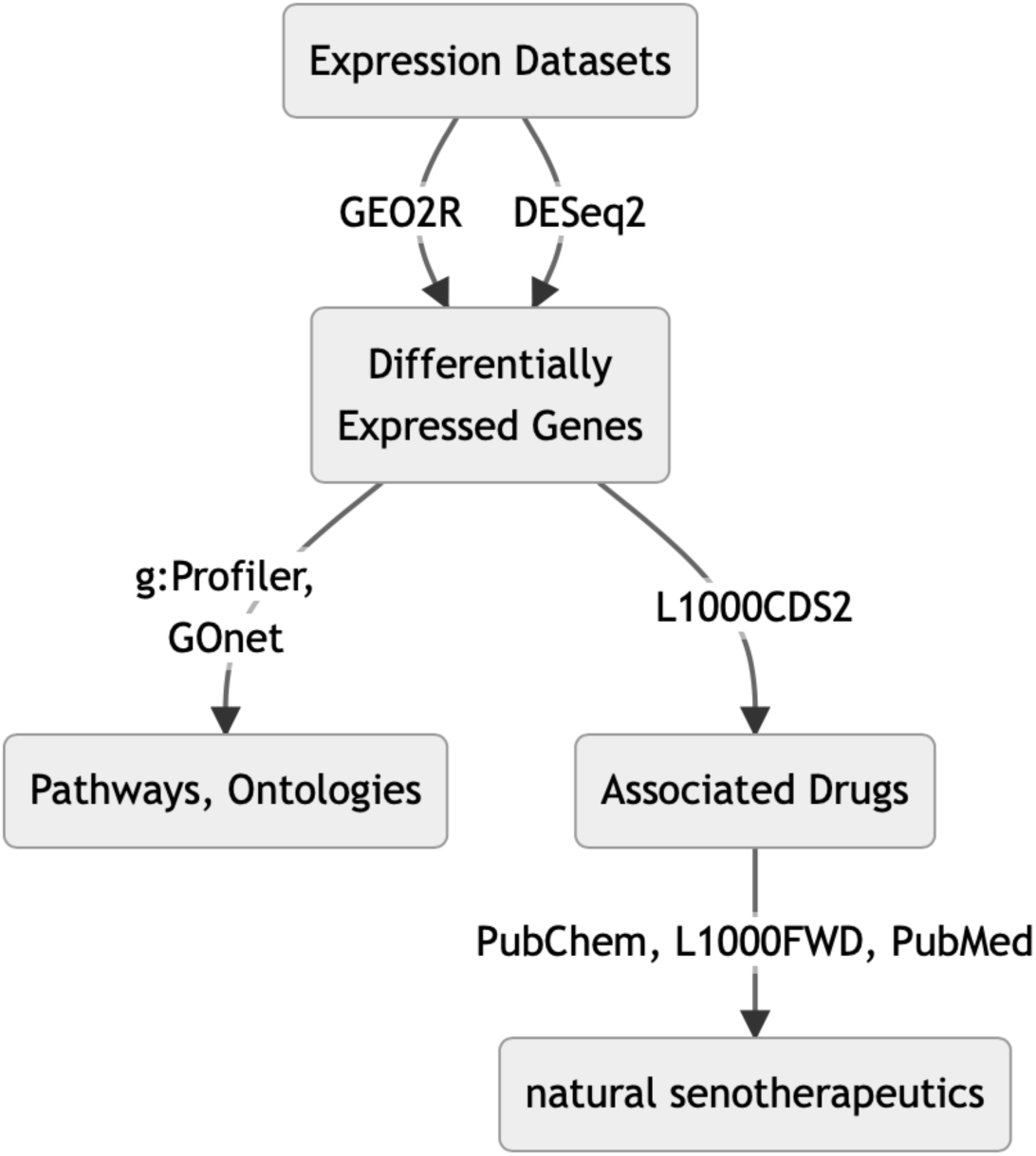
Graphical abstract showing the workflow to find natural candidate compounds with similar senolytic activity as dasatinib from expression data.

### 4.1. Expression Data

Searches for gene expression studies about drug interventions with dasatinib were conducted in the European Nucleotide Archive, the European Genome-phenome archive, the Gene Expression Omnibus (GEO) and Google Datasets (https://datasetsearch.research.google.com). Only RNA-seq and microarray datasets were considered. The search keywords included: “aging”, “senescence”, “inflammation”, “cancer”, “apoptosis”, “SASP” and “senolysis”, and they were used in combination with “dasatinib”. We found nine datasets from which the following three are subject of this paper (Table 1): dataset GSE39073, a microarray dataset containing gene expression profiles of the acute myeloid leukemia (AML) cell line Kasumi-1 subjected to long-term treatments of dasatinib (Herrmann et al., 2014); GSE9633, microarray data from experiments related to dasatinib-sensitive and dasatinib-resistant prostatic cancer cell lines (D-sensitive cell lines: 22Rv, WPMY1, VCaP, MDAPCa2b, PWR1E; D-resistant cell lines: PC3, DU145, LNCaP, HPV7, HPV10, RWPE1, RWPE2, NB11, W99, DUCaP; the strength of this dataset lies in its use of more than one cell line) (X.-D. Wang et al., 2007); and PRJNA559155, with RNASeq expression data from breast cancer cell lines exposed to either dasatinib, salinomycin, or combinations of both (Bellat et al., 2020).

The remaining datasets were excluded for reasons described in the following. Drug repositioning with gene expression signatures did not result in the identification of natural substances at the chosen cutoffs (dataset GSE59357); gene expression analysis did not result in significantly differentially expressed genes (dataset GSE69395); single-cell RNA-seq experiments cannot be directly compared against pooled cell data as stored in LINCS (accession GSE161340); cells were neither exposed to dasatinib, nor sensitivity/resistance to dasatinib was assessed as part of the experiment (datasets EGAD00001001016 and GSE14746); too few RNAseq reads were obtained after quantification of single-end reads (-r) from fastq-files in mapping-based mode (i.e. salmon quant) to the human transcriptome using salmon (Patro et al., 2017) (dataset PRJNA613485).

### 4.2. Expression analysis

Differential expression analysis of the two microarray datasets was done using the web program GEO2R (accessed May 6^th^ 2021). This program relies on GEOquery (version 2.58.0) for data retrieval, and on the R package Limma (Ritchie et al., 2015) (version 3.46.0) for the assessment of differential expression. Accordingly, these methods were applied for the selected microarray datasets (accessions GSE39073 and GSE9633). In turn, raw RNAseq sequencing reads from the accession PRJNA559155 were downloaded from the NCBI Sequence Read Archive (SRA), and mapped to the human transcriptome (GENCODE release 38) using salmon v1.4.0 (Patro et al., 2017). Differential expression analysis was then performed using DESeq2 version 1.32.0 (Love et al., 2014), for the comparison between the treatment (MDA-MB-468 cells exposed to dasatinib), and control (untreated MDA-MB-468 cells) groups. The obtained sets of differentially expressed genes were then filtered according to expression fold changes and adjusted p-values as in Supplementary Table 1.

### 4.3. Functional Analyses

Gene ontology and KEGG pathway enrichments were obtained with the g:profiler webtool (Raudvere et al., 2019; accessed 2021-11-25 and 2022-05-23) and GOnet (https://tools.dice-database.org/GOnet/) was used to perform gene annotation analysis to find genes annotated with *aging* and *senescence* (Pomaznoy et al., 2018). Finally, the NCBI gene database (https://www.ncbi.nlm.nih.gov/gene/), the human gene database GeneCards (https://www.genecards.org) and literature were used to provide gene-associated annotation information. Only default parameters were used.

### 4.4. Drug repurposing

L1000CDS^2^, a webtool that processes expression-perturbation data from the L1000 resource with a method that prioritizes small-molecule signatures that either mimic or reverse an input gene expression signatures, was used for compound identification (Duan et al., 2016). When submitting “input” (up- and downregulated genes obtained from the differential expression analysis) to L1000CDS^2^, 50 predictions (of small molecules/chemicals, characterized as perturbators of gene expression in cell lines) ranked by their overlap with the “input” signature were considered as output. Each prediction (corresponding to a perturbation) comes with seven items of information, provided in a table. This includes the rank (which is based on the overlap), the overlap (a value based on the intersection length between the input DEGs and the signature DEGs divided by the effective input, i.e. the intersection-length between input genes and L1000 genes); the venn (a schematic representation of the (mimicked or reversed) overlap of the input signature and L1000 signature), the perturbation and its associated the cell line, dose, and time, the list of overlapping genes, the predicted target genes of the perturbation, and the signature of the target/hit (Duan et al., 2016). Upregulated and downregulated genes were used as input separately, ordered by descending log2 fold changes. For the accession PRJNA559155, *mimic* mode (i.e. *mimicking* the effect of the drug by reproducing the gene expression changes associated with dasatinib-sensitivity), was chosen, and for datasets GSE9633 and GSE39073 *reverse* mode (i.e. *reversing* the disease phenotype that is susceptible to dasatinib) was chosen. The resulting lists of compounds were then manually curated, looking up each compound in the PubChem database, and PubMed, to identify natural plant metabolites.

The tabulated L1000CDS^2^ results are available online via *permanent* URLs:

AML-cell line (GSE39073, reverse): https://maayanlab.cloud/L1000CDS2/#/result/628b901ab94e3c005691571e

PC-cell line (GSE9633, reverse): https://maayanlab.cloud/L1000CDS2/#/result/619bb34fd99ec600506d5e20

BC-cell line (PRJNA559155, mimic): https://maayanlab.cloud/L1000CDS2/#/result/619f82f7d99ec600506d6086

## 5. Author contributions

Conceptualization: GF, RS, IB; Data Curation: FM; Formal Analysis: FM; Investigation: FM; Methodology: GF, SS, RS, IB; Project Administration: GF, IB; Resources: FM; Software: FM, SS; Supervision: GF, IB; Validation: FM; Visualization: FM; Writing – Original Draft Preparation: FM; Writing – Review & Editing: FM, IB, GF-

## 7. Acknowledgement

We thank Axel Kowald for critical feedback on this article.

## References

Abrahams, S., Haylett, W. L., Johnson, G., Carr, J. A., & Bardien, S. (2019). Antioxidant effects of curcumin in models of neurodegeneration, aging, oxidative and nitrosative stress: A review. Neuroscience, 406, 1–21. https://doi.org/10.1016/j.neuroscience.2019.02.020

Adams, D. J., Dai, M., Pellegrino, G., Wagner, B. K., Stern, A. M., Shamji, A. F., & Schreiber, S. L. (2012). Synthesis, cellular evaluation, and mechanism of action of piperlongumine analogs. Proceedings of the National Academy of Sciences, 109(38), 15115–15120. https://doi.org/10.1073/pnas.1212802109

Agatonovic-Kustrin, S., & Morton, D. W. (2018). The Current and Potential Therapeutic Uses of Parthenolide. In Studies in Natural Products Chemistry (Vol. 58, pp. 61–91). Elsevier. https://doi.org/10.1016/B978-0-444-64056-7.00003-9

Alikhan, M. A., Jaw, J., Shochet, L. R., Robson, K. J., Ooi, J. D., Brouwer, E., Heeringa, P., Holdsworth, S. R., & Kitching, A. R. (2021). Ageing enhances cellular immunity to myeloperoxidase and experimental anti-myeloperoxidase glomerulonephritis. Rheumatology, keab682. https://doi.org/10.1093/rheumatology/keab682

Amiri, K. I., & Richmond, A. (2005). Role of nuclear factor-kappa B in melanoma. Cancer Metastasis Reviews, 24(2), 301–313. https://doi.org/10.1007/s10555-005-1579-

Baker, D. J., Childs, B. G., Durik, M., Wijers, M. E., Sieben, C. J., Zhong, J., A. Saltness R., Jeganathan, K. B., Verzosa, G. C., Pezeshki, A., Khazaie, K., Miller, J. D., & van Deursen, J. M. (2016). Naturally occurring p16Ink4a-positive cells shorten healthy lifespan. Nature, 530(7589), 184–189. https://doi.org/10.1038/nature16932

Baker, R. G., Hayden, M. S., & Ghosh, S. (2011). NF-κB, inflammation, and metabolic disease. Cell Metabolism, 13(1), 11–22. https://doi.org/10.1016/j.cmet.2010.12.008

Bascones-Martínez, A., López-Durán, M., Cano-Sánchez, J., Sánchez-Verde, L., Díez-Rodríguez, A., Aguirre-Echebarría, P., Álvarez-Fernández, E., González-Moles, M. A., Bascones-Ilundain, J., Muzio, L. L., & Campo-Trapero, J. (2012). Differences in the expression of five senescence markers in oral cancer, oral leukoplakia and control samples in humans. Oncology Letters, 3(6), 1319–1325. https://doi.org/10.3892/ol.2012.649

Bellat, V., Verchère, A., Ashe, S. A., & Law, B. (2020). Transcriptomic insight into salinomycin mechanisms in breast cancer cell lines: Synergistic effects with dasatinib and induction of estrogen receptor β. BMC Cancer, 20(1), 661. https://doi.org/10.1186/s12885-020-07134-3

Benameur, T., Soleti, R., Panaro, M. A., La Torre, M. E., Monda, V., Messina, G., & Porro, C. (2021). Curcumin as Prospective Anti-Aging Natural Compound: Focus on Brain. Molecules (Basel, Switzerland),26(16), 4794. https://doi.org/10.3390/molecules26164794

Bernard, M., Yang, B., Migneault, F., Turgeon, J., Dieudé, M., Olivier, M.-A., Cardin, G. B., El-Diwany, M., Underwood, K., Rodier, F., & Hébert, M.-J. (2020). Autophagy drives fibroblast senescence through MTORC2 regulation. Autophagy, 16(11), 2004–2016. https://doi.org/10.1080/15548627.2020.1713640

Bianchi-Frias, D., Vakar-Lopez, F., Coleman, I. M., Plymate, S. R., Reed, M. J., & Nelson, P. S. (2010). The effects of aging on the molecular and cellular composition of the prostate microenvironment. PloS One, 5(9), e12501. https://doi.org/10.1371/journal.pone.0012501

Bielak-Zmijewska, A., Grabowska, W., Ciolko, A., Bojko, A., Mosieniak, G., Bijoch, Ł., & Sikora, E. (2019). The Role of Curcumin in the Modulation of Ageing. International Journal of Molecular Sciences, 20(5), 1239. https://doi.org/10.3390/ijms20051239

Birch, J., & Gil, J. (2020). Senescence and the SASP: Many therapeutic avenues. Genes & Development, 34(23–24), 1565–1576. https://doi.org/10.1101/gad.343129.120

Blanchard, A. A., Ma, X., Dueck, K. J., Penner, C., Cooper, S. C., Mulhall, D., Murphy, L. C., Leygue, E., & Myal, Y. (2013). Claudin 1 expression in basal-like breast cancer is related to patient age. BMC Cancer, 13, 268. https://doi.org/10.1186/1471-2407-13-268

Bork, P. M., Schmitz, M. L., Kuhnt, M., Escher, C., & Heinrich, M. (1997). Sesquiterpene lactone containing Mexican Indian medicinal plants and pure sesquiterpene lactones as potent inhibitors of transcription factor NF-κB. FEBS Letters, 402(1), 85–90. https://doi.org/10.1016/S0014-5793(96)01502-5

Bou Sleiman, M., Jha, P., Houtkooper, R., Williams, R. W., Wang, X., & Auwerx, J. (2020). The Gene-Regulatory Footprint of Aging Highlights Conserved Central Regulators. Cell Reports, 32(13), 108203. https://doi.org/10.1016/j.celrep.2020.108203

Bravo, L. (1998). Polyphenols: Chemistry, dietary sources, metabolism, and nutritional significance. Nutrition Reviews, 56(11), 317–333. https://doi.org/10.1111/j.1753-4887.1998.tb01670.x

Camell, C. D., Yousefzadeh, M. J., Zhu, Y., Prata, L. G. P. L., Huggins, M. A., Pierson, M., Zhang, L., O’Kelly, R. D., Pirtskhalava, T., Xun, P., Ejima, K., Xue, A., Tripathi, U., Espindola-Netto, J. M., Giorgadze, N., Atkinson, E. J., Inman, C. L., Johnson, K. O., Cholensky, S. H., … Robbins, P. D. (2021). Senolytics reduce coronavirus-related mortality in old mice. Science, 373(6552), eabe4832. https://doi.org/10.1126/science.abe4832

Cameron, A. R., Morrison, V. L., Levin, D., Mohan, M., Forteath, C., Beall, C., McNeilly, A. D., Balfour, D. J. K., Savinko, T., Wong, A. K. F., Viollet, B., Sakamoto, K., Fagerholm, S. C., Foretz, M., Lang, C. C., & Rena, G. (2016). Anti-Inflammatory Effects of Metformin Irrespective of Diabetes Status. Circulation Research, 119(5), 652–665. https://doi.org/10.1161/CIRCRESAHA.116.308445

Campisi, J. (2000). Cancer, aging and cellular senescence. In Vivo (Athens, Greece), 14(1), 183–188.

Castellano, J. M., Mosher, K. I., Abbey, R. J., McBride, A. A., James, M. L., Berdnik, D., Shen, J. C., Zou, B., Xie, X. S., Tingle, M., Hinkson, I. V., Angst, M. S., & Wyss-Coray, T. (2017). Human umbilical cord plasma proteins revitalize hippocampal function in aged mice. Nature, 544(7651), 488–492. https://doi.org/10.1038/nature22067

Cavalcante, M. B., Saccon, T. D., Nunes, A. D. C., Kirkland, J. L., Tchkonia, T., Schneider, A., & Masternak, M. M. (2020). Dasatinib plus quercetin prevents uterine age-related dysfunction and fibrosis in mice. Aging, 12(3), 2711–2722. https://doi.org/10.18632/aging.102772

Cevenini, E., Monti, D., & Franceschi, C. (2013). Inflamm-ageing. Current Opinion in Clinical Nutrition and Metabolic Care, 16(1), 14–20. https://doi.org/10.1097/MCO.0b013e32835ada13

Chang, J., Wang, Y., Shao, L., Laberge, R.-M., Demaria, M., Campisi, J., Janakiraman, K., Sharpless, N. E., Ding, S., Feng, W., Luo, Y., Wang, X., Aykin-Burns, N., Krager, K., Ponnappan, U., Hauer-Jensen, M., Meng, A., & Zhou, D. (2016). Clearance of senescent cells by ABT263 rejuvenates aged hematopoietic stem cells in mice. Nature Medicine, 22(1), 78–83. https://doi.org/10.1038/nm.4010

Chen, M. B., Yang, A. C., Yousef, H., Lee, D., Chen, W., Schaum, N., Lehallier, B., Quake, S. R., & Wyss-Coray, T. (2020). Brain Endothelial Cells Are Exquisite Sensors of Age-Related Circulatory Cues. Cell Reports, 30(13), 4418-4432.e4. https://doi.org/10.1016/j.celrep.2020.03.012

Childs, B. G., Baker, D. J., Kirkland, J. L., Campisi, J., & Deursen, J. M. (2014). Senescence and apoptosis: Dueling or complementary cell fates? EMBO Reports, 15(11), 1139–1153. https://doi.org/10.15252/embr.201439245

Cho, S. J., Moon, J.-S., Lee, C.-M., Choi, A. M. K., & Stout-Delgado, H. W. (2017). Glucose Transporter 1-Dependent Glycolysis Is Increased during Aging-Related Lung Fibrosis, and Phloretin Inhibits Lung Fibrosis. American Journal of Respiratory Cell and Molecular Biology, 56(4), 521–531. https://doi.org/10.1165/rcmb.2016-0225OC

Choi, H. I., Choi, G. I., Kim, E. K., Choi, Y. J., Sohn, K. C., Lee, Y., Kim, C. D., Yoon, T. J., Sohn, H. J., Han, S. H., Kim, S., Lee, J. H., & Lee, Y. H. (2011). Hair greying is associated with active hair growth. The British Journal of Dermatology, 165(6), 1183–1189. https://doi.org/10.1111/j.1365-2133.2011.10625.x

Clough, E., & Barrett, T. (2016). The Gene Expression Omnibus Database. Methods in Molecular Biology (Clifton, N.J.), 1418, 93–110. https://doi.org/10.1007/978-1-4939-3578-9_5

d’Uscio, L. V., & Katusic, Z. S. (2021). Endothelium-specific deletion of amyloid-β precursor protein exacerbates endothelial dysfunction induced by aging. Aging, 13(15), 19165–19185. https://doi.org/10.18632/aging.203401

Dierckx, T., Haidar, M., Grajchen, E., Wouters, E., Vanherle, S., Loix, M., Boeykens, A., Bylemans, D., Hardonnière, K., Kerdine-Römer, S., Bogie, J. F. J., & Hendriks, J. J. A. (2021). Phloretin suppresses neuroinflammation by autophagy-mediated Nrf2 activation in macrophages. Journal of Neuroinflammation, 18(1), 148. https://doi.org/10.1186/s12974-021-02194-z

Dörr, J. R., Yu, Y., Milanovic, M., Beuster, G., Zasada, C., Däbritz, J. H. M., Lisec, J., Lenze, D., Gerhardt, A., Schleicher, K., Kratzat, S., Purfürst, B., Walenta, S., Mueller-Klieser, W., Gräler, M., Hummel, M., Keller, U., Buck, A. K., Dörken, B., … Schmitt, C. A. (2013). Synthetic lethal metabolic targeting of cellular senescence in cancer therapy. Nature, 501(7467), 421–425. https://doi.org/10.1038/nature12437

Drewnowski, A., & Gomez-Carneros, C. (2000). Bitter taste, phytonutrients, and the consumer: A review. The American Journal of Clinical Nutrition, 72(6), 1424–1435. https://doi.org/10.1093/ajcn/72.6.1424

Duan, Q., Reid, S. P., Clark, N. R., Wang, Z., Fernandez, N. F., Rouillard, A. D., Readhead, B., Tritsch, S. R., Hodos, R., Hafner, M., Niepel, M., Sorger, P. K., Dudley, J. T., Bavari, S., Panchal, R. G., & Ma’ayan, A. (2016). L1000CDS2: LINCS L1000 characteristic direction signatures search engine. NPJ Systems Biology and Applications, 2. https://doi.org/10.1038/npjsba.2016.15

Dunkelberger, J. R., & Song, W.-C. (2010). Complement and its role in innate and adaptive immune responses. Cell Research, 20(1), 34–50. https://doi.org/10.1038/cr.2009.139

Edwards, D. N., & Bix, G. J. (2019). The Inflammatory Response After Ischemic Stroke: Targeting β2 and β1 Integrins. Frontiers in Neuroscience, 13, 540. https://doi.org/10.3389/fnins.2019.00540

El-Baba, C., Baassiri, A., Kiriako, G., Dia, B., Fadlallah, S., Moodad, S., & Darwiche, N. (2021). Terpenoids’ anti-cancer effects: Focus on autophagy. Apoptosis, 26(9–10), 491–511. https://doi.org/10.1007/s10495-021-01684-y

Everett, J. R. (2015). Academic drug discovery: Current status and prospects. Expert Opinion on Drug Discovery, 10(9), 937–944. https://doi.org/10.1517/17460441.2015.1059816

Farr, J. N., Xu, M., Weivoda, M. M., Monroe, D. G., Fraser, D. G., Onken, J. L., Negley, B. A., Sfeir, J. G., Ogrodnik, M. B., Hachfeld, C. M., LeBrasseur, N. K., Drake, M. T., Pignolo, R. J., Pirtskhalava, T., Tchkonia, T., Oursler, M. J., Kirkland, J. L., & Khosla, S. (2017). Targeting cellular senescence prevents age-related bone loss in mice. Nature Medicine, 23(9), 1072–1079. https://doi.org/10.1038/nm.4385

Franceschi, C., & Campisi, J. (2014). Chronic inflammation (inflammaging) and its potential contribution to age-associated diseases. The Journals of Gerontology. Series A, Biological Sciences and Medical Sciences, 69 Suppl 1, S4–9. https://doi.org/10.1093/gerona/glu057

Franzin, R., Stasi, A., Fiorentino, M., Stallone, G., Cantaluppi, V., Gesualdo, L., & Castellano, G. (2020). Inflammaging and Complement System: A Link Between Acute Kidney Injury and Chronic Graft Damage. Frontiers in Immunology, 11, 734. https://doi.org/10.3389/fimmu.2020.00734

Fuellen, G., Jansen, L., Cohen, A. A., Luyten, W., Gogol, M., Simm, A., Saul, N., Cirulli, F., Berry, A., Antal, P., Köhling, R., Wouters, B., & Möller, S. (2019). Health and Aging: Unifying Concepts, Scores, Biomarkers and Pathways. Aging and Disease, 10(4), 883. https://doi.org/10.14336/AD.2018.1030

Fyhrquist, F., Saijonmaa, O., & Strandberg, T. (2013). The roles of senescence and telomere shortening in cardiovascular disease. Nature Reviews Cardiology, 10(5), 274–283. https://doi.org/10.1038/nrcardio.2013.30

Gali-Muhtasib, H., Hmadi, R., Kareh, M., Tohme, R., & Darwiche, N. (2015). Cell death mechanisms of plant-derived anticancer drugs: Beyond apoptosis. Apoptosis, 20(12), 1531–1562. https://doi.org/10.1007/s10495-015-1169-2

García-García, V. A., Alameda, J. P., Page, A., & Casanova, M. L. (2021). Role of NF-κB in Ageing and Age-Related Diseases: Lessons from Genetically Modified Mouse Models. Cells, 10(8), 1906. https://doi.org/10.3390/cells10081906

Gasek, N. S., Kuchel, G. A., Kirkland, J. L., & Xu, M. (2021). Strategies for targeting senescent cells in human disease. Nature Aging, 1(10), 870–879. https://doi.org/10.1038/s43587-021-00121-8

Ghantous, A., Sinjab, A., Herceg, Z., & Darwiche, N. (2013). Parthenolide: From plant shoots to cancer roots. Drug Discovery Today, 18(17–18), 894–905. https://doi.org/10.1016/j.drudis.2013.05.005

Ghosal, G., & Chen, J. (2013). DNA damage tolerance: A double-edged sword guarding the genome. Translational Cancer Research, 2(3), 107–129. https://doi.org/10.3978/j.issn.2218-676X.2013.04.01

Goldschneider, D., & Mehlen, P. (2010). Dependence receptors: A new paradigm in cell signaling and cancer therapy. Oncogene, 29(13), 1865–1882. https://doi.org/10.1038/onc.2010.13

Gonçalves, S., Yin, K., Ito, Y., Chan, A., Olan, I., Gough, S., Cassidy, L., Serrao, E., Smith, S., Young, A., Narita, M., & Hoare, M. (2021). COX2 regulates senescence secretome composition and senescence surveillance through PGE2. Cell Reports, 34(11), 108860. https://doi.org/10.1016/j.celrep.2021.108860

Guzman, M. L., Rossi, R. M., Karnischky, L., Li, X., Peterson, D. R., Howard, D. S., & Jordan, C. T. (2005). The sesquiterpene lactone parthenolide induces apoptosis of human acute myelogenous leukemia stem and progenitor cells. Blood, 105(11), 4163–4169. https://doi.org/10.1182/blood-2004-10-4135

Hassan, F., Rehman, M. S., Khan, M. S., Ali, M. A., Javed, A., Nawaz, A., & Yang, C. (2019). Curcumin as an Alternative Epigenetic Modulator: Mechanism of Action and Potential Effects. Frontiers in Genetics, 10, 514. https://doi.org/10.3389/fgene.2019.00514

Hassane, D. C., Guzman, M. L., Corbett, C., Li, X., Abboud, R., Young, F., Liesveld, J. L., Carroll, M., & Jordan, C. T. (2008). Discovery of agents that eradicate leukemia stem cells using an in silico screen of public gene expression data. Blood, 111(12), 5654–5662. https://doi.org/10.1182/blood-2007-11-126003

Hatcher, H., Planalp, R., Cho, J., Torti, F. M., & Torti, S. V. (2008). Curcumin: From ancient medicine to current clinical trials. Cellular and Molecular Life Sciences, 65(11), 1631–1652. https://doi.org/10.1007/s00018-008-7452-4

Hayflick, L., & Moorhead, P. S. (1961). The serial cultivation of human diploid cell strains. Experimental Cell Research, 25(3), 585–621. https://doi.org/10.1016/0014-4827(61)90192-6

Herrmann, M. D., Lennerz, J. K., Bullinger, L., Bartholomae, S., Holzmann, K., Westhoff, M.-A., Corbacioglu, S., & Debatin, K.-M. (2014). Transitory dasatinib-resistant states in KIT(mut) t(8;21) acute myeloid leukemia cells correlate with altered KIT expression. Experimental Hematology, 42(2), 90–100. https://doi.org/10.1016/j.exphem.2013.10.006

Huang, H.-Y., Wu, W.-R., Wang, Y.-H., Wang, J.-W., Fang, F.-M., Tsai, J.-W., Li, S.-H., Hung, H.-C., Yu, S.-C., Lan, J., Shiue, Y.-L., Hsing, C.-H., Chen, L.-T., & Li, C.-F. (2013). ASS1 as a novel tumor suppressor gene in myxofibrosarcomas: Aberrant loss via epigenetic DNA methylation confers aggressive phenotypes, negative prognostic impact, and therapeutic relevance. Clinical Cancer Research: An Official Journal of the American Association for Cancer Research, 19(11), 2861–2872. https://doi.org/10.1158/1078-0432.CCR-12-2641

Jarada, T. N., Rokne, J. G., & Alhajj, R. (2020). A review of computational drug repositioning: Strategies, approaches, opportunities, challenges, and directions. Journal of Cheminformatics, 12(1), 46. https://doi.org/10.1186/s13321-020-00450-7

Jiang, Z.-X., Wang, Y.-N., Li, Z.-Y., Dai, Z.-H., He, Y., Chu, K., Gu, J.-Y., Ji, Y.-X., Sun, N.-X., Yang, F., & Li, W. (2021). Correction: The m6A mRNA demethylase FTO in granulosa cells retards FOS-dependent ovarian aging. Cell Death & Disease, 12(12), 1114. https://doi.org/10.1038/s41419-021-04194-6

Justice, J. N., Nambiar, A. M., Tchkonia, T., LeBrasseur, N. K., Pascual, R., Hashmi, S. K., Prata, L., Masternak, M. M., Kritchevsky, S. B., Musi, N., & Kirkland, J. L. (2019). Senolytics in idiopathic pulmonary fibrosis: Results from a first-in-human, open-label, pilot study. EBioMedicine, 40, 554–563. https://doi.org/10.1016/j.ebiom.2018.12.052

Katakura, Y., Udono, M., Katsuki, K., Nishide, H., Tabira, Y., Ikei, T., Yamashita, M., Fujiki, T., & Shirahata, S. (2009). Protein kinase C delta plays a key role in cellular senescence programs of human normal diploid cells. Journal of Biochemistry, 146(1), 87–93. https://doi.org/10.1093/jb/mvp046

Katoh, M. (2018). Multi-layered prevention and treatment of chronic inflammation, organ fibrosis and cancer associated with canonical WNT/β-catenin signaling activation (Review). International Journal of Molecular Medicine, 42(2), 713–725. https://doi.org/10.3892/ijmm.2018.3689

Kelly, C. E., Thymiakou, E., Dixon, J. E., Tanaka, S., Godwin, J., & Episkopou, V. (2013). Rnf165/Ark2C enhances BMP-Smad signaling to mediate motor axon extension. PLoS Biology, 11(4), e1001538. https://doi.org/10.1371/journal.pbio.1001538

Kim, Y. H., Choi, Y. W., Lee, J., Soh, E. Y., Kim, J.-H., & Park, T. J. (2017). Senescent tumor cells lead the collective invasion in thyroid cancer. Nature Communications, 8(1), 15208. https://doi.org/10.1038/ncomms15208

Kim, Y. R., Eom, J. I., Kim, S. J., Jeung, H. K., Cheong, J.-W., Kim, J. S., & Min, Y. H. (2010). Myeloperoxidase expression as a potential determinant of parthenolide-induced apoptosis in leukemia bulk and leukemia stem cells. The Journal of Pharmacology and Experimental Therapeutics, 335(2), 389–400. https://doi.org/10.1124/jpet.110.169367

Kirkland, J. L., & Tchkonia, T. (2020). Senolytic drugs: From discovery to translation. Journal of Internal Medicine, 288(5), 518–536. https://doi.org/10.1111/joim.13141

Knight, D. W. (1995). Feverfew: Chemistry and biological activity. Natural Product Reports, 12(3), 271–276. https://doi.org/10.1039/np9951200271

Koleti, A., Terryn, R., Stathias, V., Chung, C., Cooper, D. J., Turner, J. P., Vidović, D., Forlin, M., Kelley, T. T., D’Urso, A., Allen, B. K., Torre, D., Jagodnik, K. M., Wang, L., Jenkins, S. L., Mader, C., Niu, W., Fazel, M., Mahi, N., … Schürer, S. C. (2018). Data Portal for the Library of Integrated Network-based Cellular Signatures (LINCS) program: Integrated access to diverse large-scale cellular perturbation response data. Nucleic Acids Research, 46(D1), D558–D566. https://doi.org/10.1093/nar/gkx1063

Kubonishi, I., Machida, K., Niiya, K., Sonobe, H., Ohtsuki, Y., Iwata, K., & Miyoshi, I. (1984). Establishment of a new peroxidase-positive human myeloid cell line, PL-21. Blood, 63(2), 254–259.

Kwok, B. H., Koh, B., Ndubuisi, M. I., Elofsson, M., & Crews, C. M. (2001). The anti-inflammatory natural product parthenolide from the medicinal herb Feverfew directly binds to and inhibits IkappaB kinase. Chemistry & Biology, 8(8), 759–766. https://doi.org/10.1016/s1074-5521(01)00049-7

Lee, H. J., Lee, J., Jeong, P., Choi, J., Baek, J., Ahn, S. J., Moon, Y., Heo, J. D., Choi, Y. H., Chin, Y.-W., Kim, Y.-C., & Han, S.-Y. (2018). Discovery of a FLT3 inhibitor LDD1937 as an anti-leukemic agent for acute myeloid leukemia. Oncotarget, 9(1), 924–936. https://doi.org/10.18632/oncotarget.23221

Lehmann, M., Korfei, M., Mutze, K., Klee, S., Skronska-Wasek, W., Alsafadi, H. N., Ota, C., Costa, R., Schiller, H. B., Lindner, M., Wagner, D. E., Günther, A., & Königshoff, M. (2017). Senolytic drugs target alveolar epithelial cell function and attenuate experimental lung fibrosis ex vivo. The European Respiratory Journal, 50(2), 1602367. https://doi.org/10.1183/13993003.02367-2016

Li, Y., Zhao, Y., Qiu, C., Yang, Y., Liao, G., Wu, X., Zhang, X., Zhang, Q., Zhang, R., & Wang, Z. (2020). Role of eotaxin-1/CCL11 in sepsis-induced myocardial injury in elderly patients. Aging, 12(5), 4463–4473. https://doi.org/10.18632/aging.102896

Liao, K., Xia, B., Zhuang, Q.-Y., Hou, M.-J., Zhang, Y.-J., Luo, B., Qiu, Y., Gao, Y.-F., Li, X.-J., Chen, H.-F., Ling, W.-H., He, C.-Y., Huang, Y.-J., Lin, Y.-C., & Lin, Z.-N. (2015). Parthenolide Inhibits Cancer Stem-Like Side Population of Nasopharyngeal Carcinoma Cells via Suppression of the NF-κB/COX-2 Pathway. Theranostics, 5(3), 302–321. https://doi.org/10.7150/thno.8387

Lima, W. G., Alves-Nascimento, L. A., Andrade, J. T., Vieira, L., de Azambuja Ribeiro, R. I. M., Thomé, R. G., dos Santos, H. B., Ferreira, J. M. S., & Soares, A. C. (2019). Are the Statins promising antifungal agents against invasive candidiasis? Biomedicine & Pharmacotherapy, 111, 270–281. https://doi.org/10.1016/j.biopha.2018.12.076

Liszewski, M. K., Kolev, M., Le Friec, G., Leung, M., Bertram, P. G., Fara, A. F., Subias, M., Pickering, M. C., Drouet, C., Meri, S., Arstila, T. P., Pekkarinen, P. T., Ma, M., Cope, A., Reinheckel, T., Rodriguez de Cordoba, S., Afzali, B., Atkinson, J. P., & Kemper, C. (2013). Intracellular Complement Activation Sustains T Cell Homeostasis and Mediates Effector Differentiation. Immunity, 39(6), 1143–1157. https://doi.org/10.1016/j.immuni.2013.10.018

Liu, J., Liu, W., Lu, Y., Tian, H., Duan, C., Lu, L., Gao, G., Wu, X., Wang, X., & Yang, H. (2018). Piperlongumine restores the balance of autophagy and apoptosis by increasing BCL2 phosphorylation in rotenone-induced Parkinson disease models. Autophagy, 14(5), 845–861. https://doi.org/10.1080/15548627.2017.1390636

López-Otín, C., Blasco, M. A., Partridge, L., Serrano, M., & Kroemer, G. (2013). The Hallmarks of Aging. Cell, 153(6), 1194–1217. https://doi.org/10.1016/j.cell.2013.05.039

Love, M. I., Huber, W., & Anders, S. (2014). Moderated estimation of fold change and dispersion for RNA-seq data with DESeq2. Genome Biology, 15(12), 550. https://doi.org/10.1186/s13059-014-0550-8

Ma, L., Lu, H., Chen, R., Wu, M., Jin, Y., Zhang, J., & Wang, S. (2020). Identification of Key Genes and Potential New Biomarkers for Ovarian Aging: A Study Based on RNA-Sequencing Data. Frontiers in Genetics, 11, 590660. https://doi.org/10.3389/fgene.2020.590660

Martel, J., Ojcius, D. M., Wu, C., Peng, H., Voisin, L., Perfettini, J., Ko, Y., & Young, J. D. (2020). Emerging use of senolytics and senomorphics against aging and chronic diseases. Medicinal Research Reviews, 40(6), 2114–2131. https://doi.org/10.1002/med.21702

Menon, V. P., & Sudheer, A. R. (2007). Antioxidant and anti-inflammatory properties of curcumin. Advances in Experimental Medicine and Biology, 595, 105–125. https://doi.org/10.1007/978-0-387-46401-5_3

Morgan, H. J., Benketah, A., Olivero, C., Rees, E., Ziaj, S., Mukhtar, A., Lanfredini, S., & Patel, G. K. (2020). Hair follicle differentiation-specific keratin expression in human basal cell carcinoma. Clinical and Experimental Dermatology, 45(4), 417–425. https://doi.org/10.1111/ced.14113

Mueller, A., Schäfer, B. W., Ferrari, S., Weibel, M., Makek, M., Höchli, M., & Heizmann, C. W. (2005). The Calcium-binding Protein S100A2 Interacts with p53 and Modulates Its Transcriptional Activity. Journal of Biological Chemistry, 280(32), 29186–29193. https://doi.org/10.1074/jbc.M505000200

Nho, R. S., Peterson, M., Hergert, P., & Henke, C. A. (2013). FoxO3a (Forkhead Box O3a) Deficiency Protects Idiopathic Pulmonary Fibrosis (IPF) Fibroblasts from Type I Polymerized Collagen Matrix-Induced Apoptosis via Caveolin-1 (cav-1) and Fas. PLoS ONE, 8(4), e61017. https://doi.org/10.1371/journal.pone.0061017

Niedernhofer, L. J., & Robbins, P. D. (2018). Senotherapeutics for healthy ageing. Nature Reviews Drug Discovery, 17(5), 377–377. https://doi.org/10.1038/nrd.2018.44

Novais, E. J., Tran, V. A., Johnston, S. N., Darris, K. R., Roupas, A. J., Sessions, G. A., Shapiro, I. M., Diekman, B. O., & Risbud, M. V. (2021). Long-term treatment with senolytic drugs Dasatinib and Quercetin ameliorates age-dependent intervertebral disc degeneration in mice. Nature Communications, 12(1), 5213. https://doi.org/10.1038/s41467-021-25453-2

Ogrodnik, M., Miwa, S., Tchkonia, T., Tiniakos, D., Wilson, C. L., Lahat, A., Day, C. P., Burt, A., Palmer, A., Anstee, Q. M., Grellscheid, S. N., Hoeijmakers, J. H. J., Barnhoorn, S., Mann, D. A., Bird, T. G., Vermeij, W. P., Kirkland, J. L., Passos, J. F., von Zglinicki, T., & Jurk, D. (2017). Cellular senescence drives age-dependent hepatic steatosis. Nature Communications, 8(1), 15691. https://doi.org/10.1038/ncomms15691

Pal, S., & Tyler, J. K. (2016). Epigenetics and aging. Science Advances, 2(7), e1600584. https://doi.org/10.1126/sciadv.1600584

Paluda, A., Middleton, A. J., Rossig, C., Mace, P. D., & Day, C. L. (2022). Ubiquitin and a charged loop regulate the ubiquitin E3 ligase activity of Ark2C. Nature Communications, 13(1), 1181. https://doi.org/10.1038/s41467-022-28782-y

Panwar, P., Hedtke, T., Heinz, A., Andrault, P.-M., Hoehenwarter, W., Granville, D. J., Schmelzer, C. E. H., & Brömme, D. (2020). Expression of elastolytic cathepsins in human skin and their involvement in age-dependent elastin degradation. Biochimica Et Biophysica Acta. General Subjects, 1864(5), 129544. https://doi.org/10.1016/j.bbagen.2020.129544

Park, J.-Y., Sohn, H.-Y., Koh, Y. H., & Jo, C. (2021). Curcumin activates Nrf2 through PKCδ-mediated p62 phosphorylation at Ser351. Scientific Reports, 11(1), 8430. https://doi.org/10.1038/s41598-021-87225-8

Patro, R., Duggal, G., Love, M. I., Irizarry, R. A., & Kingsford, C. (2017). Salmon provides fast and bias-aware quantification of transcript expression. Nature Methods, 14(4), 417–419. https://doi.org/10.1038/nmeth.4197

Polosukhina, D., Singh, K., Asim, M., Barry, D. P., Allaman, M. M., Hardbower, D. M., Piazuelo, M. B., Washington, M. K., Gobert, A. P., Wilson, K. T., & Coburn, L. A. (2021). CCL11 exacerbates colitis and inflammation-associated colon tumorigenesis. Oncogene, 40(47), 6540–6546. https://doi.org/10.1038/s41388-021-02046-3

Pomaznoy, M., Ha, B., & Peters, B. (2018). GOnet: A tool for interactive Gene Ontology analysis. BMC Bioinformatics, 19(1), 470. https://doi.org/10.1186/s12859-018-2533-3

Prasad, K. N. (2016). Simultaneous activation of Nrf2 and elevation of antioxidant compounds for reducing oxidative stress and chronic inflammation in human Alzheimer’s disease. Mechanisms of Ageing and Development, 153, 41–47. https://doi.org/10.1016/j.mad.2016.01.002

Quentmeier, H., Dirks, W. G., Macleod, R. A. F., Reinhardt, J., Zaborski, M., & Drexler, H. G. (2004). Expression of HOX genes in acute leukemia cell lines with and without MLL translocations. Leukemia & Lymphoma, 45(3), 567–574. https://doi.org/10.1080/10428190310001609942

Rao, C. V. (2007). REGULATION OF COX AND LOX BY CURCUMIN. In B. B. Aggarwal, Y.-J. Surh, & S. Shishodia (Eds.), The Molecular Targets and Therapeutic Uses of Curcumin in Health and Disease (Vol. 595, pp. 213–226). Springer US. https://doi.org/10.1007/978-0-387-46401-5_9

Raudvere, U., Kolberg, L., Kuzmin, I., Arak, T., Adler, P., Peterson, H., & Vilo, J. (2019). g:Profiler: A web server for functional enrichment analysis and conversions of gene lists (2019 update). Nucleic Acids Research, 47(W1), W191–W198. https://doi.org/10.1093/nar/gkz369

Ren, L., Zhan, P., Wang, Q., Wang, C., Liu, Y., Yu, Z., & Zhang, S. (2019). Curcumin upregulates the Nrf2 system by repressing inflammatory signaling-mediated Keap1 expression in insulin-resistant conditions. Biochemical and Biophysical Research Communications, 514(3), 691–698. https://doi.org/10.1016/j.bbrc.2019.05.010

Ritchie, M. E., Phipson, B., Wu, D., Hu, Y., Law, C. W., Shi, W., & Smyth, G. K. (2015). Limma powers differential expression analyses for RNA-sequencing and microarray studies. Nucleic Acids Research, 43(7), e47–e47. https://doi.org/10.1093/nar/gkv007

Rocha, L. R., Nguyen Huu, V. A., Palomino La Torre, C., Xu, Q., Jabari, M., Krawczyk, M., Weinreb, R. N., & Skowronska-Krawczyk, D. (2020). Early removal of senescent cells protects retinal ganglion cells loss in experimental ocular hypertension. Aging Cell, 19(2). https://doi.org/10.1111/acel.13089

Romashkan, S., Chang, H., & Hadley, E. C. (2021). National Institute on Aging Workshop: Repurposing Drugs or Dietary Supplements for Their Senolytic or Senomorphic Effects: Considerations for Clinical Trials. The Journals of Gerontology: Series A, 76(6), 1144–1152. https://doi.org/10.1093/gerona/glab028

Rudin, C. M., Hann, C. L., Garon, E. B., Ribeiro de Oliveira, M., Bonomi, P. D., Camidge, D. R., Chu, Q., Giaccone, G., Khaira, D., Ramalingam, S. S., Ranson, M. R., Dive, C., McKeegan, E. M., Chyla, B. J., Dowell, B. L., Chakravartty, A., Nolan, C. E., Rudersdorf, N., Busman, T. A., … Gandhi, L. (2012). Phase II study of single-agent navitoclax (ABT-263) and biomarker correlates in patients with relapsed small cell lung cancer. Clinical Cancer Research: An Official Journal of the American Association for Cancer Research, 18(11), 3163–3169. https://doi.org/10.1158/1078-0432.CCR-11-3090

Saccon, T. D., Nagpal, R., Yadav, H., Cavalcante, M. B., Nunes, A. D. de C., Schneider, A., Gesing, A., Hughes, B., Yousefzadeh, M., Tchkonia, T., Kirkland, J. L., Niedernhofer, L. J., Robbins, P. D., & Masternak, M. M. (2021). Senolytic combination of Dasatinib and Quercetin alleviates intestinal senescence and inflammation and modulates the gut microbiome in aged mice. The Journals of Gerontology. Series A, Biological Sciences and Medical Sciences, glab002. https://doi.org/10.1093/gerona/glab002

Sage, J., De Quéral, D., Leblanc-Noblesse, E., Kurfurst, R., Schnebert, S., Perrier, E., Nizard, C., Lalmanach, G., & Lecaille, F. (2014). Differential expression of cathepsins K, S and V between young and aged Caucasian women skin epidermis. Matrix Biology: Journal of the International Society for Matrix Biology, 33, 41–46. https://doi.org/10.1016/j.matbio.2013.07.002

Salminen, A., Huuskonen, J., Ojala, J., Kauppinen, A., Kaarniranta, K., & Suuronen, T. (2008). Activation of innate immunity system during aging: NF-kB signaling is the molecular culprit of inflamm-aging. Ageing Research Reviews, 7(2), 83–105. https://doi.org/10.1016/j.arr.2007.09.002

Schafer, M. J., White, T. A., Iijima, K., Haak, A. J., Ligresti, G., Atkinson, E. J., Oberg, A. L., Birch, J., Salmonowicz, H., Zhu, Y., Mazula, D. L., Brooks, R. W., Fuhrmann-Stroissnigg, H., Pirtskhalava, T., Prakash, Y. S., Tchkonia, T., Robbins, P. D., Aubry, M. C., Passos, J. F., … LeBrasseur, N. K. (2017). Cellular senescence mediates fibrotic pulmonary disease. Nature Communications, 8, 14532. https://doi.org/10.1038/ncomms14532

Schmitt, V., Rink, L., & Uciechowski, P. (2013). The Th17/Treg balance is disturbed during aging. Experimental Gerontology, 48(12), 1379–1386. https://doi.org/10.1016/j.exger.2013.09.003

Selman, M., & Pardo, A. (2021). Fibroageing: An ageing pathological feature driven by dysregulated extracellular matrix-cell mechanobiology. Ageing Research Reviews, 70, 101393. https://doi.org/10.1016/j.arr.2021.101393

Shao, W., & Espenshade, P. J. (2012). Expanding roles for SREBP in metabolism. Cell Metabolism, 16(4), 414–419. https://doi.org/10.1016/j.cmet.2012.09.002

Sheng, S., Carey, J., Seftor, E. A., Dias, L., Hendrix, M. J., & Sager, R. (1996). Maspin acts at the cell membrane to inhibit invasion and motility of mammary and prostatic cancer cells. Proceedings of the National Academy of Sciences, 93(21), 11669–11674. https://doi.org/10.1073/pnas.93.21.11669

Shi, H., Enriquez, A., Rapadas, M., Martin, E. M. M. A., Wang, R., Moreau, J., Lim, C. K., Szot, J. O., Ip, E., Hughes, J. N., Sugimoto, K., Humphreys, D. T., McInerney-Leo, A. M., Leo, P. J., Maghzal, G. J., Halliday, J., Smith, J., Colley, A., Mark, P. R., … Dunwoodie, S. L. (2017). NAD Deficiency, Congenital Malformations, and Niacin Supplementation. The New England Journal of Medicine, 377(6), 544–552. https://doi.org/10.1056/NEJMoa1616361

Singh, A., Rangasamy, T., Thimmulappa, R. K., Lee, H., Osburn, W. O., Brigelius-Flohé, R., Kensler, T. W., Yamamoto, M., & Biswal, S. (2006). Glutathione peroxidase 2, the major cigarette smoke-inducible isoform of GPX in lungs, is regulated by Nrf2. American Journal of Respiratory Cell and Molecular Biology, 35(6), 639–650. https://doi.org/10.1165/rcmb.2005-0325OC

Son, T. G., Zou, Y., Yu, B. P., Lee, J., & Chung, H. Y. (2005). Aging effect on myeloperoxidase in rat kidney and its modulation by calorie restriction. Free Radical Research, 39(3), 283–289. https://doi.org/10.1080/10715760500053461

Spagnuolo, P. A., Hurren, R., Gronda, M., MacLean, N., Datti, A., Basheer, A., Lin, F.-H., Wang, X., Wrana, J., & Schimmer, A. D. (2013). Inhibition of intracellular dipeptidyl peptidases 8 and 9 enhances parthenolide’s anti-leukemic activity. Leukemia, 27(6), 1236–1244. https://doi.org/10.1038/leu.2013.9

Srivastava, R. K., Tang, S.-N., Zhu, W., Meeker, D., & Shankar, S. (2011). Sulforaphane synergizes with quercetin to inhibit self-renewal capacity of pancreatic cancer stem cells. Frontiers in Bioscience (Elite Edition), 3, 515–528. https://doi.org/10.2741/e266

Subramanian, A., Narayan, R., Corsello, S. M., Peck, D. D., Natoli, T. E., Lu, X., Gould, J., Davis, J. F., Tubelli, A. A., Asiedu, J. K., Lahr, D. L., Hirschman, J. E., Liu, Z., Donahue, M., Julian, B., Khan, M., Wadden, D., Smith, I. C., Lam, D., … Golub, T. R. (2017). A Next Generation Connectivity Map: L1000 Platform and the First 1,000,000 Profiles. Cell, 171(6), 1437-1452.e17. https://doi.org/10.1016/j.cell.2017.10.049

Sun, Z., Pan, X., Zou, Z., Ding, Q., Wu, G., & Peng, G. (2015). Increased SHP-1 expression results in radioresistance, inhibition of cellular senescence, and cell cycle redistribution in nasopharyngeal carcinoma cells. Radiation Oncology, 10(1), 152. https://doi.org/10.1186/s13014-015-0445-1

Suryanarayana, P., Satyanarayana, A., Balakrishna, N., Kumar, P. U., & Reddy, G. B. (2007). Effect of turmeric and curcumin on oxidative stress and antioxidant enzymes in streptozotocin-induced diabetic rat. Medical Science Monitor: International Medical Journal of Experimental and Clinical Research, 13(12), BR286–292.

Sutphin, G. L., Backer, G., Sheehan, S., Bean, S., Corban, C., Liu, T., Peters, M. J., van Meurs, J. B. J., Murabito, J. M., Johnson, A. D., Korstanje, R., & the Cohorts for Heart and Aging Research in Genomic Epidemiology (CHARGE) Consortium Gene Expression Working Group. (2017). Caenorhabditis elegansorthologs of human genes differentially expressed with age are enriched for determinants of longevity. Aging Cell, 16(4), 672–682. https://doi.org/10.1111/acel.12595

Tan, M., Heizmann, C. W., Guan, K., Schafer, B. W., & Sun, Y. (1999). Transcriptional activation of the human S100A2 promoter by wild-type p53. FEBS Letters, 445(2–3), 265–268. https://doi.org/10.1016/S0014-5793(99)00135-0

Tchkonia, T., Zhu, Y., van Deursen, J., Campisi, J., & Kirkland, J. L. (2013). Cellular senescence and the senescent secretory phenotype: Therapeutic opportunities. Journal of Clinical Investigation, 123(3), 966–972. https://doi.org/10.1172/JCI64098

Thongsom, S., Suginta, W., Lee, K. J., Choe, H., & Talabnin, C. (2017). Piperlongumine induces G2/M phase arrest and apoptosis in cholangiocarcinoma cells through the ROS-JNK-ERK signaling pathway. Apoptosis: An International Journal on Programmed Cell Death, 22(11), 1473–1484. https://doi.org/10.1007/s10495-017-1422-y

Ukraintseva, S., Duan, M., Arbeev, K., Wu, D., Bagley, O., Yashkin, A. P., Gorbunova, G., Akushevich, I., Kulminski, A., & Yashin, A. (2021). Interactions Between Genes From Aging Pathways May Influence Human Lifespan and Improve Animal to Human Translation. Frontiers in Cell and Developmental Biology, 9, 692020. https://doi.org/10.3389/fcell.2021.692020

van den HEUVEL, A. P. J., Schulze, A., & Burgering, B. M. T. (2005). Direct control of caveolin-1 expression by FOXO transcription factors. Biochemical Journal, 385(3), 795–802. https://doi.org/10.1042/BJ20041449

Wakabayashi, K., Isozaki, T., Tsubokura, Y., Fukuse, S., & Kasama, T. (2021). Eotaxin-1/CCL11 is involved in cell migration in rheumatoid arthritis. Scientific Reports, 11(1), 7937. https://doi.org/10.1038/s41598-021-87199-7

Wang, C.-H., Wang, L.-K., Wu, C.-C., Chen, M.-L., Kuo, C.-Y., Shyu, R.-Y., & Tsai, F.-M. (2020). Cathepsin V Mediates the Tazarotene-induced Gene 1-induced Reduction in Invasion in Colorectal Cancer Cells. Cell Biochemistry and Biophysics, 78(4), 483–494. https://doi.org/10.1007/s12013-020-00940-3

Wang, H., Wang, Z., Huang, Y., Zhou, Y., Sheng, X., Jiang, Q., Wang, Y., Luo, P., Luo, M., & Shi, C. (2020). Senolytics (DQ) Mitigates Radiation Ulcers by Removing Senescent Cells. Frontiers in Oncology, 9, 1576. https://doi.org/10.3389/fonc.2019.01576

Wang, X.-D., Reeves, K., Luo, F. R., Xu, L.-A., Lee, F., Clark, E., & Huang, F. (2007). Identification of candidate predictive and surrogate molecular markers for dasatinib in prostate cancer: Rationale for patient selection and efficacy monitoring. Genome Biology, 8(11), R255. https://doi.org/10.1186/gb-2007-811-r255

Wang, Y., Chang, J., Liu, X., Zhang, X., Zhang, S., Zhang, X., Zhou, D., & Zheng, G. (2016). Discovery of piperlongumine as a potential novel lead for the development of senolytic agents. Aging, 8(11), 2915–2926. https://doi.org/10.18632/aging.101100

Weisberg, E., Liu, Q., Nelson, E., Kung, A. L., Christie, A. L., Bronson, R., Sattler, M., Sanda, T., Zhao, Z., Hur, W., Mitsiades, C., Smith, R., Daley, J. F., Stone, R., Galinsky, I., Griffin, J. D., & Gray, N. (2012). Using combination therapy to override stromal-mediated chemoresistance in mutant FLT3-positive AML: Synergism between FLT3 inhibitors, dasatinib/multi-targeted inhibitors and JAK inhibitors. Leukemia, 26(10), 2233–2244. https://doi.org/10.1038/leu.2012.96

Whisler, R. L., Chen, M., Beiqing, L., & Carle, K. W. (1997). Impaired induction of c-fos/c-jun genes and of transcriptional regulatory proteins binding distinct c-fos/c-jun promoter elements in activated human T cells during aging. Cellular Immunology, 175(1), 41–50. https://doi.org/10.1006/cimm.1996.1048

Wiley, C. D., & Campisi, J. (2021). The metabolic roots of senescence: Mechanisms and opportunities for intervention. Nature Metabolism, 3(10), 1290–1301. https://doi.org/10.1038/s42255-021-00483-8

Wills-Karp, M. (2007). Complement activation pathways: A bridge between innate and adaptive immune responses in asthma. Proceedings of the American Thoracic Society, 4(3), 247–251. https://doi.org/10.1513/pats.200704-046AW

Wynn, T. A., & Ramalingam, T. R. (2012). Mechanisms of fibrosis: Therapeutic translation for fibrotic disease. Nature Medicine, 18(7), 1028–1040. https://doi.org/10.1038/nm.2807

Xu, M., Pirtskhalava, T., Farr, J. N., Weigand, B. M., Palmer, A. K., Weivoda, M. M., Inman, C. L., Ogrodnik, M. B., Hachfeld, C. M., Fraser, D. G., Onken, J. L., Johnson, K. O., Verzosa, G. C., Langhi, L. G. P., Weigl, M., Giorgadze, N., LeBrasseur, N. K., Miller, J. D., Jurk, D., … Kirkland, J. L. (2018). Senolytics improve physical function and increase lifespan in old age. Nature Medicine, 24(8), 1246–1256. https://doi.org/10.1038/s41591-018-0092-9

Xu, Q., Long, Q., Zhu, D., Fu, D., Zhang, B., Han, L., Qian, M., Guo, J., Xu, J., Cao, L., Chin, Y. E., Coppé, J.-P., Lam, E. W.-F., Campisi, J., & Sun, Y. (2019). Targeting amphiregulin (AREG) derived from senescent stromal cells diminishes cancer resistance and averts programmed cell death 1 ligand (PD-L1)-mediated immunosuppression. Aging Cell, 18(6), e13027. https://doi.org/10.1111/acel.13027

Xue, H., Li, J., Xie, H., & Wang, Y. (2018). Review of Drug Repositioning Approaches and Resources. International Journal of Biological Sciences, 14(10), 1232–1244. https://doi.org/10.7150/ijbs.24612

Ying, Y., Jiang, C., Zhang, M., Jin, J., Ge, S., & Wang, X. (2019). Phloretin protects against cardiac damage and remodeling via restoring SIRT1 and anti-inflammatory effects in the streptozotocin-induced diabetic mouse model. Aging, 11(9), 2822–2835. https://doi.org/10.18632/aging.101954

Zaman, M. S., Barman, S. K., Corley, S. M., Wilkins, M. R., Malladi, C. S., & Wu, M. J. (2021). Transcriptomic insights into the zinc homeostasis of MCF-7 breast cancer cells via next-generation RNA sequencing. Metallomics, 13(6), mfab026. https://doi.org/10.1093/mtomcs/mfab026

Zhang, L., Zhao, J., Mu, X., McGowan, S. J., Angelini, L., O’Kelly, R. D., Yousefzadeh, M. J., Sakamoto, A., Aversa, Z., LeBrasseur, N. K., Suh, Y., Huard, J., Kamenecka, T. M., Niedernhofer, L. J., & Robbins, P. D. (2021). Novel small molecule inhibition of IKK/NF-κB activation reduces markers of senescence and improves healthspan in mouse models of aging. Aging Cell, 20(12), e13486. https://doi.org/10.1111/acel.13486

Zhang, X., Zhang, S., Liu, X., Wang, Y., Chang, J., Zhang, X., Mackintosh, S. G., Tackett, A. J., He, Y., Lv, D., Laberge, R.-M., Campisi, J., Wang, J., Zheng, G., & Zhou, D. (2018). Oxidation resistance 1 is a novel senolytic target. Aging Cell, 17(4), e12780. https://doi.org/10.1111/acel.12780

Zhou, Y., Xin, X., Wang, L., Wang, B., Chen, L., Liu, O., Rowe, D. W., & Xu, M. (2021). Senolytics improve bone forming potential of bone marrow mesenchymal stem cells from aged mice. Npj Regenerative Medicine, 6(1), 34. https://doi.org/10.1038/s41536-021-00145-z

Zhu, M., Meng, P., Ling, X., & Zhou, L. (2020). Advancements in therapeutic drugs targeting of senescence. Therapeutic Advances in Chronic Disease, 11, 204062232096412. https://doi.org/10.1177/2040622320964125

Zhu, Y., Doornebal, E. J., Pirtskhalava, T., Giorgadze, N., Wentworth, M., Fuhrmann-Stroissnigg, H., Niedernhofer, L. J., Robbins, P. D., Tchkonia, T., & Kirkland, J. L. (2017). New agents that target senescent cells: The flavone, fisetin, and the BCL-XL inhibitors, A1331852 and A1155463. Aging, 9(3), 955–963. https://doi.org/10.18632/aging.101202

Zhu, Y., Tchkonia, T., Pirtskhalava, T., Gower, A. C., Ding, H., Giorgadze, N., Palmer, A. K., Ikeno, Y., Hubbard, G. B., Lenburg, M., O’Hara, S. P., LaRusso, N. F., Miller, J. D., Roos, C. M., Verzosa, G. C., LeBrasseur, N. K., Wren, J. D., Farr, J. N., Khosla, S., … Kirkland, J. L. (2015b). The Achilles’ heel of senescent cells: From transcriptome to senolytic drugs. Aging Cell, 14(4), 644–658. https://doi.org/10.1111/acel.12344

Zia, A., Farkhondeh, T., Pourbagher-Shahri, A. M., & Samarghandian, S. (2021). The role of curcumin in aging and senescence: Molecular mechanisms. Biomedicine & Pharmacotherapy, 134, 111119. https://doi.org/10.1016/j.biopha.2020.111119

