## Supplementary material for "Computational identification of natural senotherapeutic compounds that mimic dasatinib based on gene expression data": Suppl Material (Tables, Figure, Texts)

Reference is made herein to the supplementary excel-sheet **dasatinib_compid_V03 Supplem_Material.xlsx**

**Supplementary Tables**

ST1: Number of upregulated/ downregulated genes of each of the datasets

| Dataset | Adjusted p-value | Upregulated DEGs | Downregulated DEGs |
| --- | --- | --- | --- |
| GSE39073, LFC = 2 | < 0.01 | 190 | 192 |
| GSE9633 LFC = 2 | < 0.05 | 138 | 51 |
| PJRNA559155 LFC = 1.5 | < 0.05 | 109 | 80 |

ST2: Genes annotated with the biological process *aging.* PRKCD was annotated also with the biological process *cellular* *senescence*; DEGs from AML-dataset GSE39073 were used. Genes were identified with the GOnet-webtool (<https://tools.dice-database.org/GOnet/>) with analysis type “GO-term annotation”. LFC: log2 fold change.

| Gene | LFC | association with aging | Reference |
| --- | --- | --- | --- |
| KYNU | 5.16 | Knockout in C. elegans resulted in lifespan extension of >20% | (Sutphin et al., 2017) |
| FOS | 3.58 | associated with ovarian aging, role in activated human T-cells | (Jiang et al., 2021; Whisler et al., 1997) |
| PRKCD | 3.37 | tumor suppressor protein, cell cycle regulator, apoptosis regulation. Association with senescence-induction in human diploid cells | (Katakura et al., 2009) |
| ITGB2 | 2.73 | probable role in ovarian aging, higher expression in ovaries from old mice | (Ma et al., 2020) |
| BCL2 | 2.62 | senolytic BCL2 inhibition leads to apoptosis of senescent cells, may influence human lifespan | (Ukraintseva et al., 2021; M. Zhu et al., 2020) |
| MPO | -3.03 | aging, immune cells. High MPO levels can be reduced by a calorie-restricted diet | (Son et al., 2005) |
| APP | -2.79 | protective response to aging-induced inflammation in endothelial cells | (d’Uscio & Katusic, 2021) |
| TIMP2 | -2.36 | associated with synaptic plasticity and cognition in aged mice | (Castellano et al., 2017) |

ST3: Genes annotated with the biological process *aging*, DEGs from prostate-cancer (PC) dataset GSE9633. Genes were identified with the GOnet-webtool (<https://tools.dice-database.org/GOnet/>) with analysis type “GO-term annotation”. LFC: log2 fold change.

| Gene | LFC | association with aging | Reference |
| --- | --- | --- | --- |
| SERPINB5 | 6.66 | tumor suppressor and senescence-associated biomarker | (Bascones-Martínez et al., 2012; Sheng et al., 1996) |
| CTSV | 4.06 | Matrix-degrading protease associated with skin aging | (Panwar et al., 2020) |
| CLDN1 | 4.01 | Polymorphisms are associated with age (55 or older) in breast cancer patients | (Blanchard et al., 2013; Katoh, 2018) |
| TGFBR2 | 3.61 | Regulation of cell survival and apoptosis with a potential role in human longevity | (Ukraintseva et al., 2021) |
| CDKN2A | 3.12 | senescence-associated marker | (Childs et al., 2014) |
| ASS1 | 3.06 | Association with Alzheimer’s disease | (Prasad, 2016) |

ST4: Genes annotated with the biological process *aging*, DEGs from breast-cancer dataset PRJNA559155 were used. Genes were identified with the GOnet-webtool (<https://tools.dice-database.org/GOnet/>) with analysis type “GO-term annotation”. LFC: log2 fold change.

| Gene | LFC | association with aging | Reference |
| --- | --- | --- | --- |
| CCL11 | -9.78 | SASP factor aging- and inflammation associated plasma cytokine | (Cameron et al., 2016) |
| KRT25 | -7.64 | hair greying | (Choi et al., 2011) |
| RNF165 | 3.38 | enhances BMP-smad signalling and mediates motor axon extension, knockout associated with premature death in mice | (Kelly et al., 2013) |
| SREBF1 | 1.83 | regulates lipid homeostasis, possibly an aging-associated transcription factor | (Shao & Espenshade, 2012) (Bou Sleiman et al., 2020) |

ST5: Overlapping genes between the up- (and down)regulated genes of AML-dataset GSE39073, and down (and up)regulated genes of piperlongumine-induced expression changes from the L1000 database.

| Input up/ signature down | input down/ signature up |
| --- | --- |
| ACSL1 | DDAH1 |
| ATP8B4 | FBXO21 |
| CTSG | SLC38A1 |
| EIF1AY | TSPAN13 |
| FLT3 |  |
| HCK |  |
| KDM5D |  |
| LYZ |  |
| PLAC8 |  |
| PRKCD |  |
| PTPN6 |  |
| RNASE2 |  |
| RPS6KA1 |  |
| TNFRSF10B |  |
| TNS3 |  |

ST6: Enriched biological processes associated with *apoptosis* of overlapping genes between the up- (and down)regulated genes of AML-dataset GSE39073, and down (and up)regulated genes of piperlongumine-induced expression changes from the L1000 database. Obtained from enrichr version 2021. The complete table of enriched biological processes is available in the excel-sheet: overlapGO_NOMO1_PL_AML-05-15-21

| Term | Adjusted P-value | Genes |
| --- | --- | --- |
| regulation of apoptotic process (GO:0042981) | 1.43E-02 | HCK;FLT3;RPS6KA1;TNFRSF10B;PTPN6 |
| negative regulation of apoptotic process (GO:0043066) | 2.03E-02 | HCK;PRKCD;RPS6KA1;CTSG |
| intrinsic apoptotic signaling pathway (GO:0097193) | 4.28E-02 | PRKCD;TNFRSF10B |
| regulation of glial cell apoptotic process (GO:0034350) | 4.28E-02 | PRKCD |
| TRAIL-activated apoptotic signaling pathway (GO:0036462) | 4.28E-02 | TNFRSF10B |
| modulation by symbiont of host apoptotic process (GO:0052150) | 4.28E-02 | CTSG |
| negative regulation of glial cell apoptotic process (GO:0034351) | 4.28E-02 | PRKCD |
| negative regulation by symbiont of host apoptotic process (GO:0033668) | 4.28E-02 | CTSG |
| intrinsic apoptotic signaling pathway in response to oxidative stress (GO:0008631) | 4.54E-02 | PRKCD |

ST7: Overlapping genes between the up- (and down)regulated genes of PC-dataset GSE9633, and down (and up) genes of piperlongumine-induced expression changes from the L1000 database.

| Input up/ signature down | input down/ signature up |
| --- | --- |
| AHNAK2 | LEF1 |
| ALDH1A3 |  |
| AREG |  |
| C3 |  |
| CAPG |  |
| CST6 |  |
| DDX60 |  |
| FERMT1 |  |
| ITGA3 |  |
| KRT7 |  |
| LAMA3 |  |
| RAC2 |  |
| RRAS |  |
| S100A2 |  |
| TGFBR2 |  |
| ZBED2 |  |

ST8: Enriched biological processes associated with *apoptosis* of overlapping genes between the up- (and down)regulated genes of PC-dataset GSE9633, and down (and up) genes of piperlongumine-induced expression changes from the L1000 database. Obtained from enrichr version 2021. The complete table of enriched biological processes is available in the excel-sheet; overlapGO_PL_PC3_PC-dataset

| Term | Adjusted P-value | Genes |
| --- | --- | --- |
| engulfment of apoptotic cell (GO:0043652) | 4.26E-02 | RAC2 |
| positive regulation of apoptotic cell clearance (GO:2000427) | 4.17E-02 | C3 |
| regulation of apoptotic cell clearance (GO:2000425) | 4.17E-02 | C3 |

Supplementary Figures

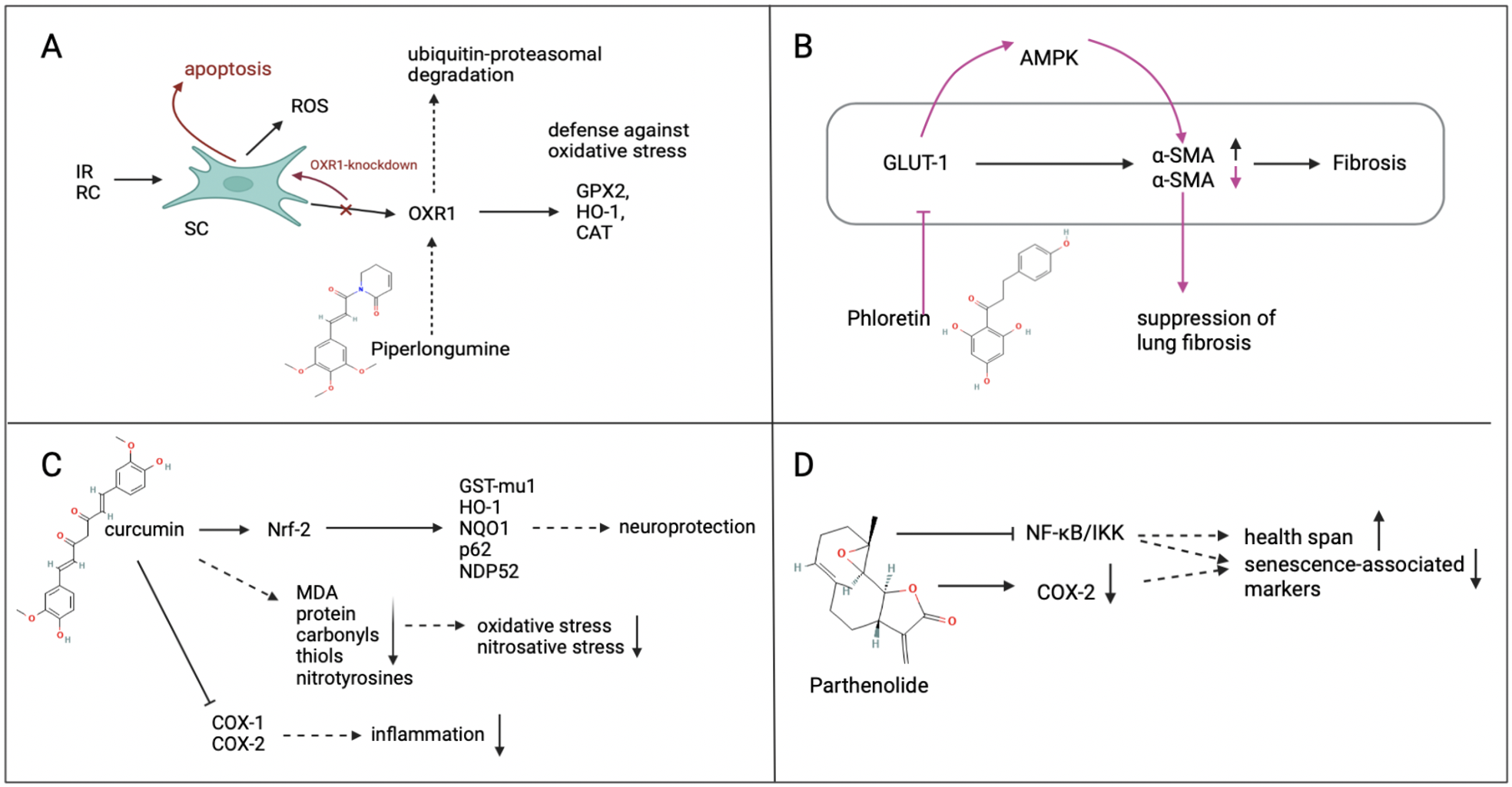

Supplementary Figure 1. Proposed models of how the identified compounds may interact with senescence/aging. (A), potential mechanisms of senolytic action of Piperlongumine. As concluded by X. Zhang et al. (2018) piperlongumine directly binds to OXR1, suppressing the expression of antioxidant genes which leads to apoptosis of senescent fibroblasts. (B) Phloretin is a potent SLC2A1 (GLUT-1) inhibitor. According to experiments conducted by Cho et al., (2017), its administration leads to activation of AMP-Kinase and reduces expression of α-smooth muscle actin in lung fibroblasts leading to suppression of lung fibrosis in aged mice (Cho et al., 2017). (C) Proposed mechanism of anti-inflammatory, neuroprotective and anti-aging effects of curcumin as identified by Park et al., 2021. (D) Proposed targets of Parthenolide and potential age-related effects according to Zhang et al. (2021): Parthenolide inhibits NF-κB and IBK-Kinase (IKK)-complex, and downregulates COX-2 leading to reduction of SASP-factors and an increase in health span in mice model (L. Zhang et al., 2021). SC: senescent cell; RO: reactive oxygen species; IR: ionizing radiation; RC: replicative senescence.

Supplementary texts

2.1 Genes annotated with the gene ontology term *aging* in the Kasumi-1 AML dataset

(for genes FOS, PRKCD, BCL2 see main text and ST2).

-	The highly upregulated gene **KNYU** (kynureninase) encodes an enzyme involved in the synthesis of NAD cofactors (Shi et al., 2017). The proto-oncogene **FOS** (or AP-1), was previously linked to ovarian aging and was associated with activated human T-cells during aging (Jiang et al., 2021; Whisler et al., 1997).

-	The **ITGB2** gene encodes an integrin beta-chain of integrins, used by leukocytes to migrate towards areas of inflammation, e.g. following a stroke (Edwards & Bix, 2019). Previously, ITGB2 was identified as a key gene in ovarian aging (Ma et al., 2020).

-	The **MPO** gene encodes for myeloperoxidase, a major constituent of neutrophil granules. Cellular immunity to MPO develops in aged mice and may be connected to the increased incidence of Anti-Neutrophil Cytoplasmic Antibodies (ANCA)-associated vasculitis, an autoimmune disease characterized by small blood vessel inflammation which commonly affects kidneys and respiratory tract in humans (Alikhan et al., 2021). Increased MPO activity with aging may be connected to increased immune cells-recruitment, contributing to protein oxidation-accumulation during the aging process (Son et al., 2005). In rats fed an ad libitum diet, higher levels of MPO were observed with age, and effects were attenuated by a calorie restricted dietary regimen (Son et al., 2005).

-	Next, the **APP** gene encodes a cell surface receptor and transmembrane precursor protein ([NCBI Gene ID 351](https://www.ncbi.nlm.nih.gov/gene/351)). APP expression is increased in aged wild-type mouse aortas, especially in endothelial cells, probably stimulated by the presence of proinflammatory cytokines (d’Uscio & Katusic, 2021). The upregulation of APP in endothelial cells represents and adaptive/ protective response to aging-induced inflammation, and in turn, depletion of APP worsens endothelial dysfunction induced by aging (Chen et al., 2020; d’Uscio & Katusic, 2021).

-	**TIMP2** is a natural matrix metalloproteinase inhibitor and able to directly suppress endothelial cell-proliferation ([NCBI Gene ID 7077](https://www.ncbi.nlm.nih.gov/gene/7077)). TIMP2 appeared in the brain of aged mice after systemic administration of human umbilical cord plasma proteins, given to mice with the intention to increase synaptic plasticity and hippocampal-dependent cognition (Castellano et al., 2017). The group found that TIMP2 was necessary for the cognitive benefits conferred by cord plasma which promoted plasticity and reduced brain-aging (Castellano et al., 2017)

### 2.2 Genes annotated with the gene ontology term *aging* in dasatinib-sensitive prostatic cancer cell lines (for genes SERPINB5, TGFBR2 & CDKN2A see main text and ST3.)

-	The gene **CTSV** encodes cathepsin V, a potent elastase and matrix-degrading protease that degrades elastin fibers; decreased expression is associated with skin aging (Panwar et al., 2020; Sage et al., 2014). In cancer, CTSV expression increases the expression of activated urokinase-type plasminogen activator (uPA), a protein associated with epithelial-mesenchymal transition (EMT), along with the number of migrating and invading cells (C.-H. Wang et al., 2020).

-	Next, **CLDN1** is a tight junction protein and target gene of beta-catenin, a signaling pathway involved in myofibroblast activation and subsequent fibrosis (Katoh, 2018). There is also an association with CLDN1 expression and age in breast cancer patients aged 55 and older (Blanchard et al., 2013).

-	Further, the **ASS1** gene was identified, which catalyzes the penultimate step of arginine synthesis, it is a tumor suppressor in myxofibrosarcoma and loss of expression was linked to more aggressive tumor progression; in Alzheimer’s disease (AD) defective expression of ASS1 peptides initiate and promote AD (Huang et al., 2013; Prasad, 2016).

### 2.3 Genes annotated with the gene ontology term *aging* in the MDA-MB-468 breast cancer (BC) dataset (for the gene CCL11, see main text and ST4.)

-	The second-most downregulated gene was **KRT25**, encoding a hair follicle specific keratin. KRT25 is increased in the hair follicles of graying hair, and higher expressed in white hair vs. black hair (Choi et al., 2011), it is further expressed in human basal cell carcinoma in the inner root sheath (Morgan et al., 2020).

-	In turn, the gene **RNF165**, upregulated in response to dasatinib-treatment, is a RING E3 ligase and positive regulator of TGF-beta signaling (Kelly et al., 2013) as it mediates the attachment of degradative ubiquitin chains to negative regulators of the TGF-beta pathway, leading to destruction (Paluda et al., 2022). RNF165 is only expressed in the nervous system and enhances BMP-Smad signaling to mediate motor axon extension (Kelly et al., 2013), but was also found to be implicated in zinc homeostasis in MCF-7 breast cancer cells (Zaman et al., 2021). Loss of RNF165 in mice leads to motor innervation defects, resulting in death and wasting before weaning (Kelly et al., 2013).

-	**SBREF1** is involved in regulating lipid homeostasis, possibly having wide-ranging effects in the aging process in conditions such as type II diabetes and cancer (Shao & Espenshade, 2012), and is thus a putative aging-associated transcription factor (Bou Sleiman et al., 2020).

2.4 Supplementary information on other natural compounds found by our approach.

**Parthenolide** is a sesquiterpene lactone, referred to as the “medieval aspirin” because of its longterm use to treat headache, fever and arthritis (El-Baba et al., 2021; Knight, 1995). The small molecule is mainly isolated from the medicinal herb feverfew (*Tanacetum parthenium*) but is found in other plants from the daisy family, e.g. in the German Chamomile (Agatonovic-Kustrin & Morton, 2018; Kwok et al., 2001). Parthenolide is an NF-κB pathway inhibitor at noncytotoxic pharmacological concentrations (Ghantous et al., 2013). And, unlike most NF-κB pathway inhibitors, parthenolide does not have antioxidant properties based on its chemical structure (Bork et al., 1997), but inhibits NF-κB by directly binding to NF-κB subunits (Ghantous et al., 2013). Parthenolide also inhibits the IKB kinase complex, which phosphorylates two NF-κB inhibitors: IκBα (IKKα) and IκBβ (IKKβ) resulting in proteasomal degradation (Ghantous et al., 2013). IKKβ plays a role in cytokine-mediated signaling (Kwok et al., 2001), and inhibition of IKK/NF-κB activation reduced inflammatory markers and improved health in an aging mouse model (L. Zhang et al., 2021). A promising feature of parthenolide is the selectivity against cancer stem cells by inhibiting NF-κB, activating TP53 and inducing oxidative stress and apoptosis, and inhibiting tumor cell differentiation in primary AML cells (Guzman et al., 2005; Hassane et al., 2008; Kim et al., 2010; Spagnuolo et al., 2013). Parthenolide also targets COX2, a gene that is upregulated in senescence and regulates SASP component composition (Gonçalves et al., 2021; Liao et al., 2015).

**Curcumin** is a polyphenolic compound isolated from the rhizome of the perennial herb turmeric (*Curcuma longa*) (Bielak-Zmijewska et al., 2019) with potential senotherapeuic properties. Generally, curcumin has anti-inflammatory, anti-oxidative and antineoplastic activity, along with an ability to inhibit tumor cell proliferation and chemically induced carcinogenesis (Zia et al., 2021). The compound affects a range of biological pathways, including the NRF-2 and NF-kB pathway (Hassan et al., 2019; Hatcher et al., 2008). It decreases the activity of proinflammatory cyclooxygenase COX-2 and, via suppression of NF-kB, reduces NF-kB-regulated genes, and SASP factors, such as CXCR-4, proinflammatory interleukins IL1A, IL6, CXCL8; and TNF, while activating the protective stress responsive factor NRF-2 (Birch & Gil, 2020; Hassan et al., 2019; Y. H. Kim et al., 2017; Rao, 2007).

Curcumin is also an epigenetic modulator, and has been shown to reduce histone acetylation, prevent histone de-acetylation, upregulate the expression of tumor-suppressive miRNAs and to reduce the expression of oncogenic miRNAs (Hassan et al., 2019). Epigenetic changes during aging and replicative senescence include a reduction of bulk levels of core histones, a change in patterns of histone modification and DNA methylation leading to altered local accessibility of DNA, which would otherwise result in (unwanted) aberrant gene expression and genomic instability (Pal & Tyler, 2016).

In a model of diabetic mice, treatment of epithelial progenitor cells with curcumin reduced the number of SA-β-gal-positive senescent cells, promoted blood flow and upregulated the pro-angiogenic factors VEGFA and ANGPT1 (You et al., 2017).
Cucumin also reduced the number of SA-β-gal-positive senescent murine embryonic fibroblasts at a concentration of 5µM, a dose where quercetin was not effective (Yousefzadeh et al., 2018). Also here a killing of senescent cells could not be shown and curcumin was declared as “senotherapeutic”.

**Phloretin** is an inhibitor of glucose transporter 1 (SLC2A1). SLC2A1-dependent glycolysis has been found to be activated in a model of lung fibrosis and is suggested to regulate age-dependent fibrogenesis, shown by the inhibition of bleomycin-induced lung fibrosis *in vivo* (Cho et al., 2017). Fibrosis (or scarring) is a feature of most chronic inflammatory conditions, and involves the accumulation of ECM proteins, activated fibroblasts and inflammatory cells (Wynn & Ramalingam, 2012). The tendency to develop tissue fibrosis is linked to aging, and the ECM is proposed to be a key player fundamental to a process termed “fibroageing” (Selman & Pardo, 2021). SLC2A1 dependent glycolysis regulates the activation of fibrogenesis in aged lungs (Cho et al., 2017). Phloretin is able to inhibit SLC2A1 expression via activation of AMPK leading to suppressed production of α-smooth muscle actin in lungs of aged mice, a cytoskeletal component of activated fibroblasts (Cho et al., 2017). In vitro, phloretin inhibited high-glucose induced inflammation and fibrosis via SIRT1 expression (Ying et al., 2019). Additionally, phloretin reduced the inflammatory phenotype in macrophages by activation of NRF2 signaling which was attributed to AMPK-dependent activation of autophagy (Dierckx et al., 2021).
